# Corticotrophin-Releasing Hormone neurons of Barrington’s nucleus: Probabilistic, spinally-gated control of bladder pressure and micturition

**DOI:** 10.1101/683334

**Authors:** H. Ito, A.C. Sales, C.H. Fry, A.J. Kanai, M.J. Drake, A.E. Pickering

## Abstract

Micturition, the co-ordinated process of expulsion of urine from the bladder, requires precise control of bladder and urethral sphincter via parasympathetic, sympathetic and somatic motoneurons. In adult mammals this involves a spinobulbospinal control circuit incorporating Barrington’s nucleus in the pons (Barr). The largest Barr cell population is comprised of pontospinal glutamatergic neurons that express corticotrophin-releasing hormone. There is evidence that Barr^CRH^ neurons can generate bladder contractions but it is unknown whether they act as a simple switch or a high-fidelity pre-parasympathetic motor drive and whether their activation can actually trigger voids. Combined opto- and chemo-genetic manipulations along with recordings in mice shows that Barr^CRH^ neurons provide a probabilistic drive that generates co-ordinated voids or non-voiding contractions depending on the phase of the micturition cycle. These findings inform a new inferential model of micturition and emphasise the importance of the state of the spinal gating circuit in the generation of voiding.

## Introduction

The regulated production, storage and elimination of liquid waste as urine (micturition) plays a critical homeostatic role in maintaining the health of organisms. Like breathing, this involves precisely co-ordinated autonomic and somatic motor drives and has both voluntary and autonomous (involuntary) control mechanisms. The power of the autonomous drive is illustrated by the challenge faced by anyone *“caught short”* away from a socially acceptable location for urination. Disorders of autonomous micturition include overactive bladder syndrome, enuresis and following frontal lobe lesions. Barrington’s nucleus, also known as the pontine micturition centre, is a key site for the control of urination^1^. The prevailing concept of the neural control of micturition is that afferent information from the bladder is conveyed via the spinal cord to the periaqueductal gray (PAG) in the midbrain where it is integrated with information from higher centres such as hypothalamus and cortex^2–5^. When the bladder is full, a threshold is reached and a neural command to void is relayed from Barrington’s nucleus to the lumbosacral parasympathetic neurons and urethral sphincter motoneurons. Lesion of Barrington’s nucleus^1^ or acute transection of the pons abolishes micturition^6,7^; in contrast, supra-collicular decerebration or transection of PAG does not stop micturition in cats, rats or mice^6,8,9^. Similar studies showed electrical or chemical stimulation of Barrington’s nucleus induces bladder contraction^10–13^. Functional imaging studies in humans^14,15^ and rats^16^ found activity in the dorsal pons during voiding. Thus, Barrington’s nucleus is thought to be pivotal in the voiding reflex and is believed to be the pre-parasympathetic control centre.

The majority of Barrington’s neurons express corticotropin releasing hormone (CRH) in humans^17^ and rodents^18–21^ and CRH axons from the pons terminate in the vicinity of the sacral parasympathetic neurons^20–22^. The role of these CRH positive neurons in Barrington’s (Barr^CRH^) neurons has recently been explored using CRH^CRE^ mice to enable specific opto- and chemo-genetic manipulation of their activity^22,23^. These studies indicated that Barr^CRH^ neurons were glutamatergic, their activation caused bladder contraction and increasing their excitability increased the probability of micturition. A second smaller subgroup of oestrogen receptor type-1 positive neurons in Barrington’s (Barr^ESR1^) have been shown to be important for control of the urethral sphincter in voluntary scent marking with urine^23^. Further a group of layer-5 pyramidal neurons in the primary motor cortex plays a role in the descending control of voluntary urination via their projections to Barrington’s nucleus^24^.

Single-unit recordings of micturition-related neurons in the vicinity of Barrington’s nucleus in rats and cats showed multiple different patterns of activity with either increased or decreased firing during bladder contractions^25–29^. These results were thought to reflect the neural heterogeneity within Barrington’s and/or be due to the complex neural circuits in nearby brainstem sites involved the regulation of other pelvic visceral functions. More recent microwire recordings of the dorsal pons in rats reported neurons in the vicinity of Barrington’s nucleus that have more homogeneous firing patterns, characterised by tonic activity with phasic bursts that were temporally associated with the voiding phase of the micturition cycle^30^ but they also show bursts of activity between voids that were not associated with increases in bladder pressure. A common technical limitation of these pontine neural recordings is the difficulty of identifying specific cell populations during or after recordings. This has been addressed for populations of Barrington’s neurons through fibre-photometry of genetically encoded calcium indicators in mice^22–24^ to show that Barr^CRH^ and Barr^ESR1^ neuronal activity increases around the time of voiding / scent marking respectively. However, the limited temporal and spatial resolution of the indicator and technique limits the ability to address whether this activity drives or follows micturition behaviour and the associated increase in bladder pressure. Therefore, the exact role of the Barr^CRH^ neurons in micturition remains unclear and it is presumed that they act as a central control centre generating a pre-motor parasympathetic drive to the bladder but it is not known whether they are sufficient on their own to generate a co-ordinated void through their actions on spinal circuits.

Here, we study the role of Barr^CRH^ neurons in the autonomous micturition cycle using opto- and chemo-genetic interventions as well as recordings of the firing activity of identified Barr^CRH^ neurons *in vivo*. This has informed the development of a model indicating that these Barr^CRH^ neurons provide a probabilistic signal to spinal circuits that is gated to trigger either non-voiding bladder contractions, which enable inferences to be made about the degree of bladder fullness, or voiding if a threshold level of pressure has been reached.

## Results

### Barr^CRH^ neurons modulate micturition

Recent studies have drawn contrasting conclusions about the role and importance of Barr^CRH^ neurons in the regulation of voiding^22,23^. To further define their role, the light-activated cation channel ChR2 (channelrhodopsin-2) was selectively expressed in Barr^CRH^ neurons of CRH^CRE^ mice^31^ using a Cre-dependent adeno-associated viral vector (AAV-EF1*α*-DIO-ChR2-mcherry) (Figure 1A-C). Saline infusion to the bladder produced a regular cycle of autonomous micturition (voids at ∼5 minute intervals, average infusion rate 23±4µl/min). Tonic unilateral activation of Barr^CRH^ neurons (5-20Hz x 10ms, 465nm light pulses, applied across 3 voiding cycles) produced an increase of micturition frequency (153.2±15.6% at 10Hz, Figure 1D) manifesting as a significant shortening in inter-void interval associated with a reduction of the threshold pressure for voiding (121.4±7.1% at 10Hz, Figure 1E). Similar illumination in control mice (CRH^CRE^ mice injected with AAV-DIO-hm4Di-mCherry instead of ChR2) had no effect on voiding frequency.

**Figure 1.**
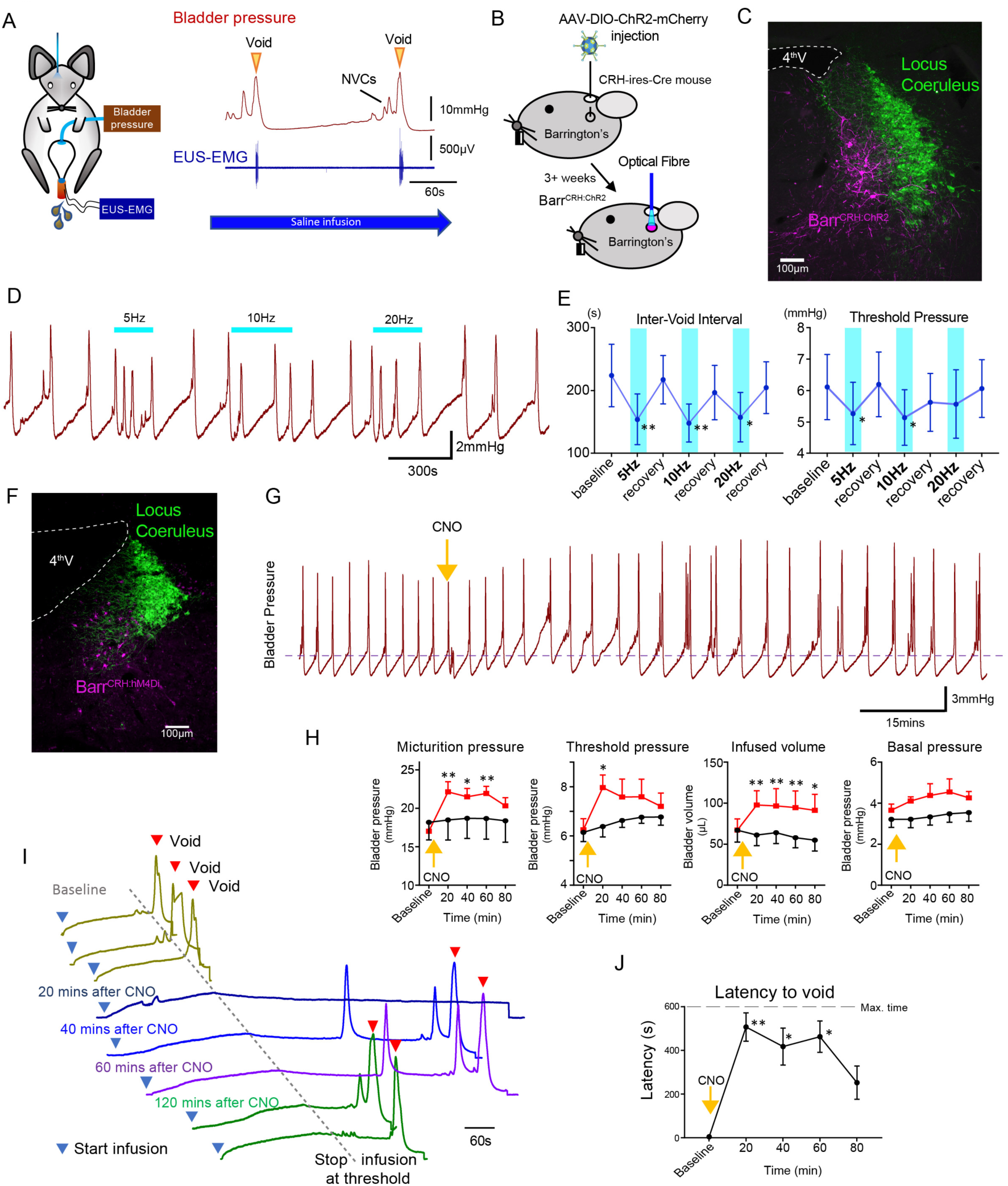
BarrCRH neurons modulate the micturition cycle. A) *in vivo* recording in urethane anaesthetized mice with optical fibre positioned above Barrington’s nucleus in the pons with bladder pressure and external urethral sphincter activity monitoring during the micturition cycle (with continuous bladder filling). B) AAV injection to dorsal pons in CRHCre mice followed after 3 weeks by opto-activation by light (465nm) from an optical fibre. C) Post-hoc histology after stereotaxic injection of AAV-DIO-ChR2-mCherry demonstrating transduction of BarrCRH neurons in sections of dorsal pons. Immunohistochemistry for mCherry (Magenta) shows transduced Barr neurons and Tyrosine hydroxylase (Green) marks the adjacent Locus Coeruleus. D) Periods of maintained unilateral opto-activation of BarrCRH:ChR2 increased the frequency of micturition (light pulsed at 5, 10 and 20Hz x 20ms for 3 micturition cycles). E) Continuous opto-activation (5 and 10Hz) reversibly shortened the inter-void interval (*n=7* mice). Similarly, the threshold for voiding was reversibly reduced with 5Hz stimulation (Friedman test with Dunn’s multiple comparisons to prior unstimulated state *P<0.05, **P<0.01). F) Transduction of BarrCRH with inhibitory DREADD using AAV-DIO-hM4Di-mCherry demonstrated with immunocytochemistry for mCherry (magenta) and Tyrosine hydroxylase (green) to mark the Locus Coeruleus laterally. G) Administration of the DREADD ligand CNO (5mg/kg, i.p) slowed the frequency of micturition seen with continuous saline infusion to the bladder. H) The chemogenetic inhibition of BarrCRH:hM4Di neurons caused an increase in the voiding threshold (129.7 ± 8.9%), volume infused before void (161.9±16.9%) and micturition pressure (131.7 ± 6.9%) compared to baseline (RM-ANOVA with Holm-Sidak’s post hoc or Friedman’s test with Dunn’s post hoc, *P<0.05, **P<0.01) versus control mice (BarrCRH:ChR2, *n=9* per group). In each case this peaked at around 20 minutes after CNO and reversed slowly. I) using an intermittent bladder infusion protocol (to a maximum bladder pressure of 15mmHg) CNO administration inhibited voiding with (J) a large increase in the latency to void – equivalent to urinary retention (*n=5*) (RM-ANOVA with Holm-Sidak’s post hoc, *P<0.05, **P<0.01).

Chemogenetic-inhibition of Barr^CRH^ neurons slowed voiding (Figure 1F, following bilateral injection of AAV-DIO-hM4Di to CRH^CRE^ mice). The administration of Clozapine N-oxide (CNO, 5mg/kg i.p) produced a prolonged reduction in the frequency of voids seen during a continuous infusion protocol (reduced to 66.8±6.3% of control at 20 mins, Figure 1G). This was associated with increases of threshold pressure, fill volume and micturition pressure (Figure 1H). The chemogenetic inhibition of Barr^CRH^ during this continuous filling protocol caused the bladder to become increasingly distended leading to incomplete voids. To control for this effect a ‘fill and hold’ protocol was employed to fill the bladder to a maximum pressure of 15mmHg (close to the threshold pressure for voiding). This volume was held for up to 10 minutes until either the mouse voided spontaneously, or the bladder was manually emptied, and the filling cycle restarted (Figure 1I). Using this protocol, chemogenetic inhibition caused a prolongation of the interval to void (from 207±17s at baseline to 738±60 s at 20mins after CNO). This effectively produced a period of urinary retention, that recovered over the space of ∼2 hours (Figure 1J).

These findings indicate that Barr^CRH^ neurons have a potent ability to modulate the autonomous micturition cycle and that their basal level of activity is of functional importance.

### Barr^CRH^ neurons do not simply act as high-fidelity controllers of bladder pressure

To assess whether Barr^CRH^ neurons act as a tightly-coupled, pre-motor drive to bladder parasympathetic neurons^2,32^ bladder pressure was recorded while parametrically opto-activating Barrington’s nucleus unilaterally (9.5±0.3mW, 465nm). In initial experiments, opto-activation of Barr^CRH^ (20ms x 20Hz for 5 s) evoked non-voiding contractions of the bladder (eNVC, Supp Figure 1A & B, with the bladder filled to half of its threshold capacity, *n=7* mice). Varying stimulus frequencies and pulse durations produced modestly graded changes in eNVC with a 20ms x 20Hz protocol producing near maximal responses (3.9±0.8 mmHg, Supp Figure 1B). The eNVC had a consistent latency to onset of 1.3±0.1s and a time to peak of 6.0±0.3s following stimulus onset and an average duration of 8.2±0.6s. With each of the stimulus parameters there were ‘failures’ where there was no detectable bladder response (Supp figure 1B, Figure 2D). The probability of eNVC increased with stimulation frequency (71.4±7.6% at 2.5Hz and 97.1±2.5% at 20Hz) with 20Hz being the most reliable. Single light pulses of longer duration (1-3 s) were also able to reliably generate eNVC. In contrast, illumination in control mice (CRH^CRE^ mice injected with AAV-DIO-hm4Di-mCherry instead of ChR2) had no effect on the bladder pressure (101.4±0.6% at 20Hz x 20ms, *n=3*).

**Figure 2.**
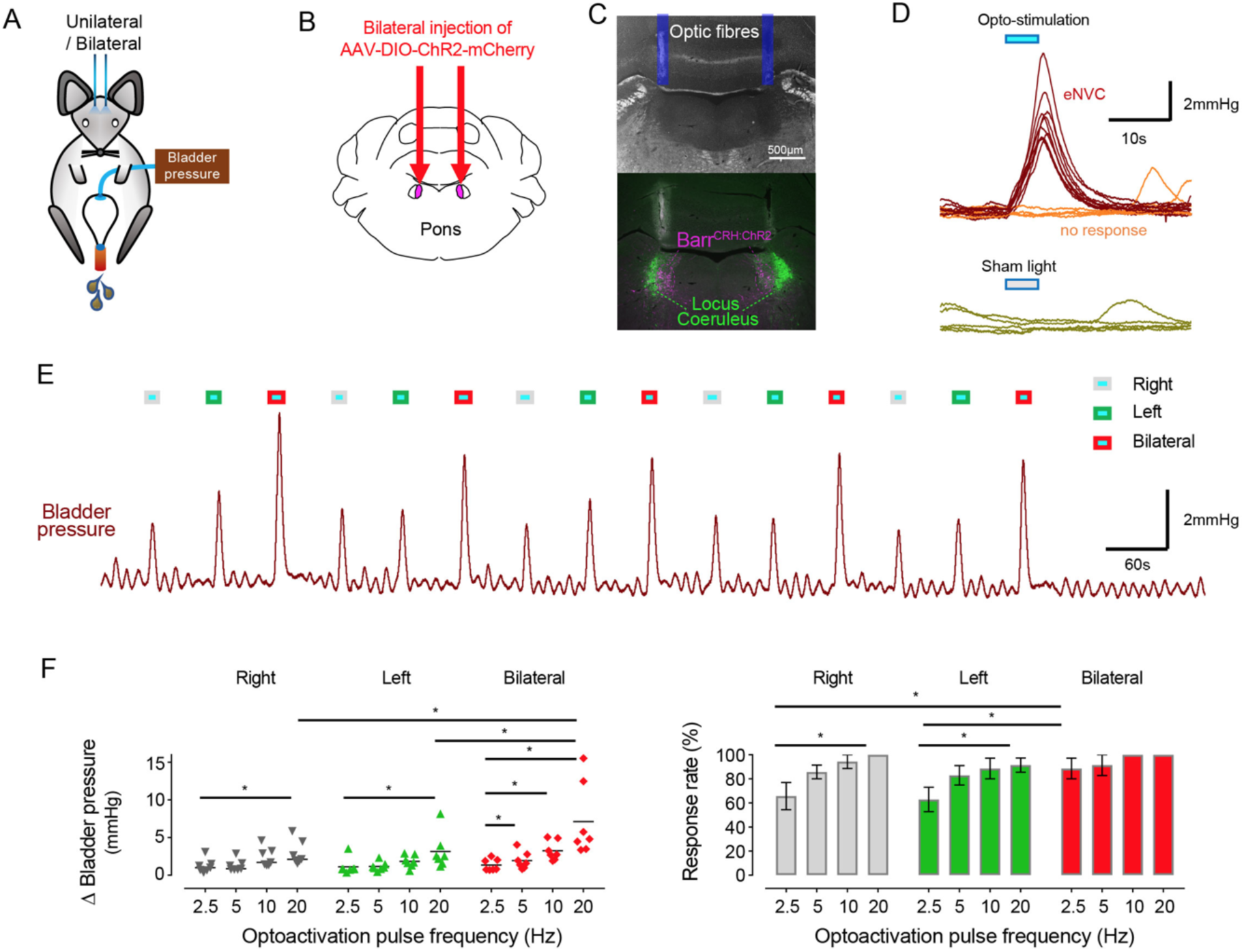
Phasic optoactivation of Barr^CRH^ evokes non-voiding contractions of the bladder. (A) Bladder pressure recordings with unilateral or bilateral opto-activation following (B) bilateral injection of AAV-DIO-ChR2-mCherry. (C) Confirmation of bilateral Barr^CRH:ChR2^ transduction and optic fibre targeting (immuno for mCherry – magenta and TH - green). (D) Phasic opto-activation (20ms x 20Hz, 5s) of Barr^CRH^ neurons evoked non-voiding contractions (eNVCs, with the bladder ∼half full, static). These eNVCs had a stereotyped shape and a relatively constant latency. In addition, there were ‘failures’ where no response was evoked by an identical stimulus. A comparison of the effects of unilateral versus bilateral stimulation (E, F) showed that bilateral stimulation evoked larger events more reliably than either side alone (each point represents data from a single mouse, *n=7* mice, Friedman’s test, *P<0.05, **P<0.01).

It was postulated that recruitment of a larger population of Barr^CRH^ neurons in synchrony might be more effective in triggering bladder contractions. This was tested with bilateral expression of ChR2 and a dual-fibre optical cannula allowing independent activation of one, the other or both Barrington’s nuclei (Figure 2). Bilateral activation of Barr^CRH^ produced larger eNVC (7.1±2.0mmHg at 20ms and 20Hz, *n=7* mice) than optoactivation of either side alone (2.9±0.5 and 3.1±0.8mmHg for right or left side, respectively), particularly at higher frequencies of stimulation (Figure 2E&F). The effect of bilateral stimulation on eNVC amplitude was additive rather than synergistic. The probability of generating eNVC was increased with bilateral stimulation (evident at lower stimulus frequencies i.e. increased by 154±18% *vs* right alone or by 158±17% *vs* left alone at 2.5Hz, Figure 2F). It was notable that, with the bladder filled to half of its threshold capacity, bilateral Barr^CRH^ stimulation never triggered voids.

Previous anatomical studies with retrograde tracing using pseudorabies virus have suggested that there is a route for communication between afferent neurons innervating the distal colon and Barrington’s nucleus and further that Barrington’s neurons were activated by distention of the distal colon^33^. This has led to the suggestion that Barrington’s nucleus may control the lower gastrointestinal tract as well as the lower urinary tract. Indeed, defaecation was suggested to occur on occasion (but not quantitated) following opto-activation of Barr^CRH^ neurons in mice implying a role in motor control^22^. To test the relationship between Barr^CRH^ and activity in the distal colon, a balloon catheter was inserted to monitor pressure. Distal colonic pressure was not synchronised with bladder pressure during the normal micturition cycle (Supp. Figure 2). Furthermore, optogenetic activation of Barr^CRH^ did not alter distal colonic pressure (Supp. Figure 2), despite the generation of eNVC, suggesting that this population of Barrington’s neurons is not involved in motor control of the colon.

These findings support the proposal that Barr^CRH^ neurons can selectively generate bladder contractions and indicate that this is a probabilistic process, with failures, rather than being a simple high-fidelity pre-motor drive to the bladder.

### Bladder pressure responses to Barr^CRH^ drive augments with progress through the micturition cycle

This raised the question of whether the stage of the micturition cycle influences the bladder pressure response to Barr^CRH^ opto-activation as the cycle phase may modulate Barr^CRH^ neuronal excitability. During continuous bladder filling, it was noted that the amplitude of Barr^CRH^ eNVC increased progressively through the micturition cycle (increase of 17.0±3.9 fold, comparing eNVC obtained during the 2^nd^ versus 5^th^ quintile of micturition cycle, Figure 3A-D). Similarly, the probability of obtaining a bladder contraction with optoactivation also increased with progressive filling, with most “failures” being seen when the bladder was <40% filled (Figure 3E). The same phase dependence of eNVC was also apparent with bilateral stimulation of Barr^CRH^ (Sup Figure 1C&D).

**Figure 3.**
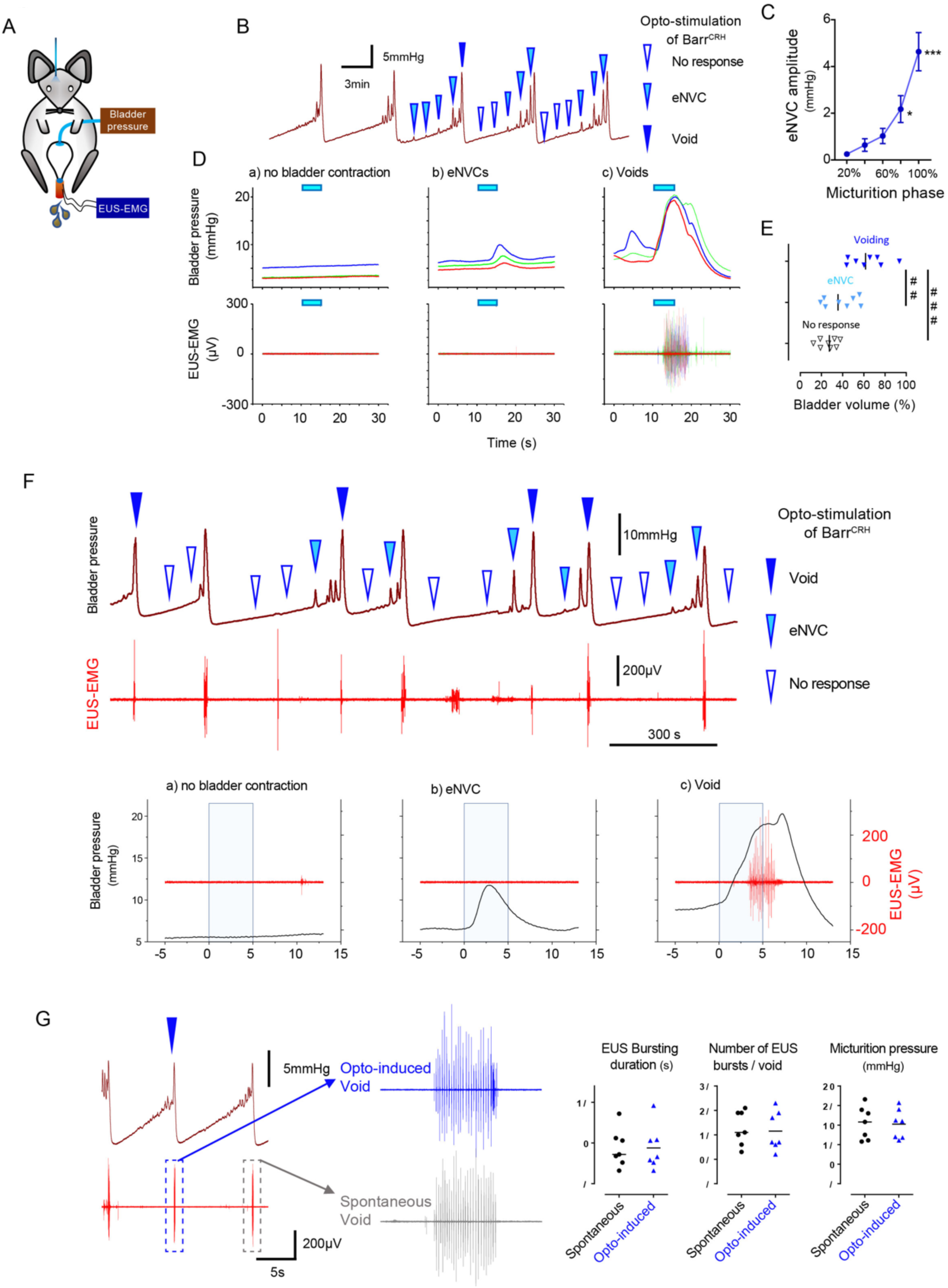
Dynamics of Barr^CRH^ evoked events through the micturition cycle. A) Experimental set-up with unilateral opto-activation of Barr^CRH^ neurons. B) Continuous infusion cystometry with episodic opto-activation (20Hz x 20ms for 5s) applied at different phases of the micturition cycle generates eNVC of incrementing amplitude as the cycle progresses. C) There is a substantial increase in the amplitude of the eNVC as the micturition cycle progresses (17.0±3.9 fold comparing eNVC from the 2^nd^ and 5^th^ quintiles of the cycle) (Repeated measures one-way ANOVA followed by Dunnet’s test). D) Overlaid bladder pressure responses to the same optogenetic stimulus applied at different phases of the cycle can trigger either no response or eNVCs or full voiding contractions that show a stereotyped morphology and latency. E) Analysis of the stage of the voiding cycle where each type of response was triggered showed that voiding contractions were significantly more likely to be evoked later in the voiding cycle (each symbol represents the average position of such events in each mouse, *n=8*) (Tukey’s test). F) Only the Barr^CRH^-evoked voids were associated with EUS activity (shown blown up below). G) The bursting pattern of EUS activity was similar with both Barr^CRH^-evoked and spontaneous voids (*n=7*).

The phase-dependence of eNVC amplitude may, in part, be a consequence of bladder distension, leading to raised passive detrusor tension and an increase of length-dependent contractions. To test this proposition, the effect of pelvic nerve stimulation was assessed in the pithed decerebrate, arterially-perfused mouse preparation^9,34^. The amplitude of bladder contractions induced by pelvic nerve stimulation (4-20Hz, 10V, 3sec) increased with bladder filling (Supp Figure 3) with a doubling (2.2±0.34 fold at 20Hz) of the pressure generated between empty bladder and 70µl fill (close to voiding threshold in an intact mouse). However, this amplitude increase plateaued at a volume of ∼50µl – and showed a much less steep relationship than that observed for Barr^CRH^ eNVC *in vivo* which increased by 17-fold over the same range of bladder distension. Additionally, this relationship did not account for the observed probabilistic nature of eNVC, as failures were never observed with pelvic nerve stimulation.

### Barr^CRH^ stimulation can conditionally trigger complete voids

Although tonic stimulation of Barr^CRH^ increased voiding frequency, it was not possible to trigger full voiding contractions with phasic Barr^CRH^ stimulation with the bladder up to 50% filled, even with bilateral stimulation. However, by applying bursts of stimulation systematically at points through the micturition cycle it was possible to trigger fully co-ordinated voids by activating Barr^CRH^ neurons later in the cycle (>50% filled, Figure 3D-G). The pattern and amplitude of the evoked bladder contraction was similar to that seen with spontaneous voids and they occurred at a similar latency to eNVC (Figure 3G). In addition, voiding was complete and the empty bladder relaxed to the basal pressure level after each void (Figure 3F).

The mouse external urethral sphincter (EUS) shows bursting activity during spontaneous voids which facilitates urine expulsion^9,23^. Injections of pseudorabies virus into either the bladder or EUS has shown labelling in the vicinity of Barrington’s nucleus, suggesting it is part of the EUS control circuit^35–37^. However, recent evidence suggests that it is the Barr^ESR-1^, rather than Barr^CRH^, neurons which project to local circuit interneurons in L4-5 that may regulate EUS motoneurons^23^ analogous to the lumbar spinal coordinating centre (LSCC ^38^). Therefore, recordings were made from the EUS to investigate the relationship of the voiding-associated bursting to Barr^CRH^ activation. The Barr^CRH^ eNVC (irrespective of their magnitude) were never associated with EUS activity (Figure 3G). However, when Barr^CRH^ activation evoked a voiding contraction then bursting EUS activity was always found (Figure 3G). These Barr^CRH^ induced voids had EUS activity that was indistinguishable from spontaneous voids in terms of burst duration, frequency and individual burst length (Figure 3 H&I).

These results indicate that the Barr^CRH^ neurons can trigger voids that are in all aspects similar to those seen spontaneously but that can be triggered to occur earlier in the normal micturition cycle.

### Spinal drive from Barr^CRH^ neurons is sufficient to generate eNVC and voids

The axons from Barr^CRH^ were noted to provide a specific innervation of the sacral parasympathetic neurons but not to the ventral horn at the level of Onuf’s nucleus (Supp Figure 4). To investigate whether optogenetic stimulation of spinal axons of Barr^CRH^ is sufficient to directly generate eNVC, bladder pressure was recorded while light was applied from an optic fibre located above the spinal cord (at T11). Optogenetic stimuli (either 20ms x 20Hz for 5 s or single 1s pulse) applied to the spinal cord reliably induced bladder contractions (Supp Figure 5, *p*=0.025, bladder half filled). These eNVCs tended to occur with a shorter latency than those evoked directly from pontine stimulation (1.0±0.2s vs 1.26±0.1s, *n=5*). Similarly, during continuous bladder filling, spinal activation could trigger full voids (Supp Figure 5D). These data support the principle that the Barr^CRH^ neurons can evoke both voiding and eNVC through their spinal projections.

**Figure 4.**
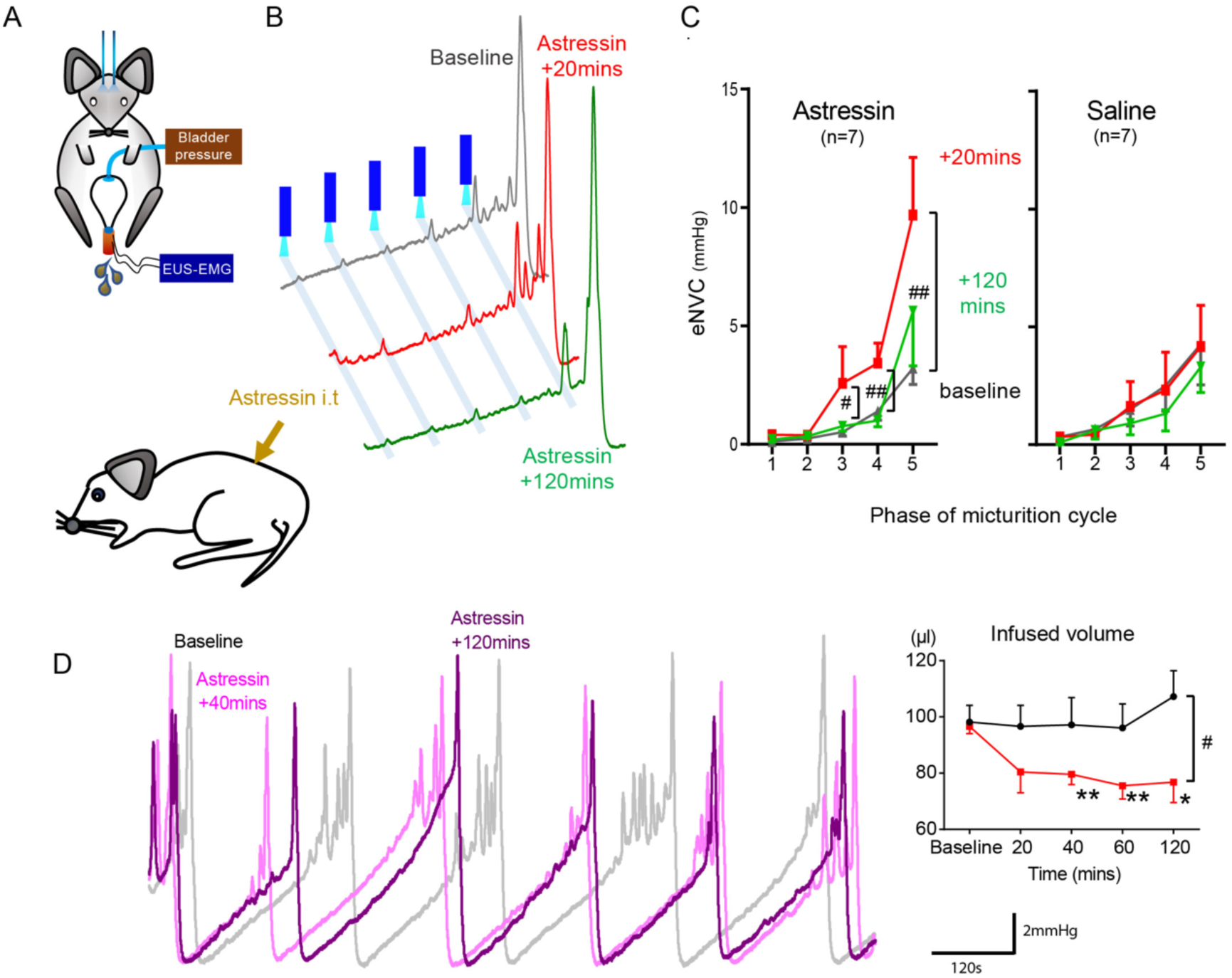
Spinal CRH inhibits the bladder response to Barr^CRH^ neuronal optoactivation. A) Assessment of the influence of intrathecal Astressin (CRH antagonist) on the bladder pressure response to bilateral optoactivation of Barrington’s. B) Intrathecal Astressin (5µg) reversibly increased the amplitude of Barr^CRH^ eNVC (*n=7* mice). C) Summary data for the action of Astressin on eNVC (versus vehicle control) showing that the augmentation of amplitude was particularly marked towards the end of the micturition cycle (Related samples Friedman’s test by ranks). D) Even without Barr^CRH^ opto-stimulation Astressin reversibly increased the frequency of voiding compared both to baseline and an intrathecal vehicle control group (*n=9*) (vs baseline with Related samples Friedman’s test by ranks and vs vehicle with Mann-Whitney U test).

**Figure 5.**
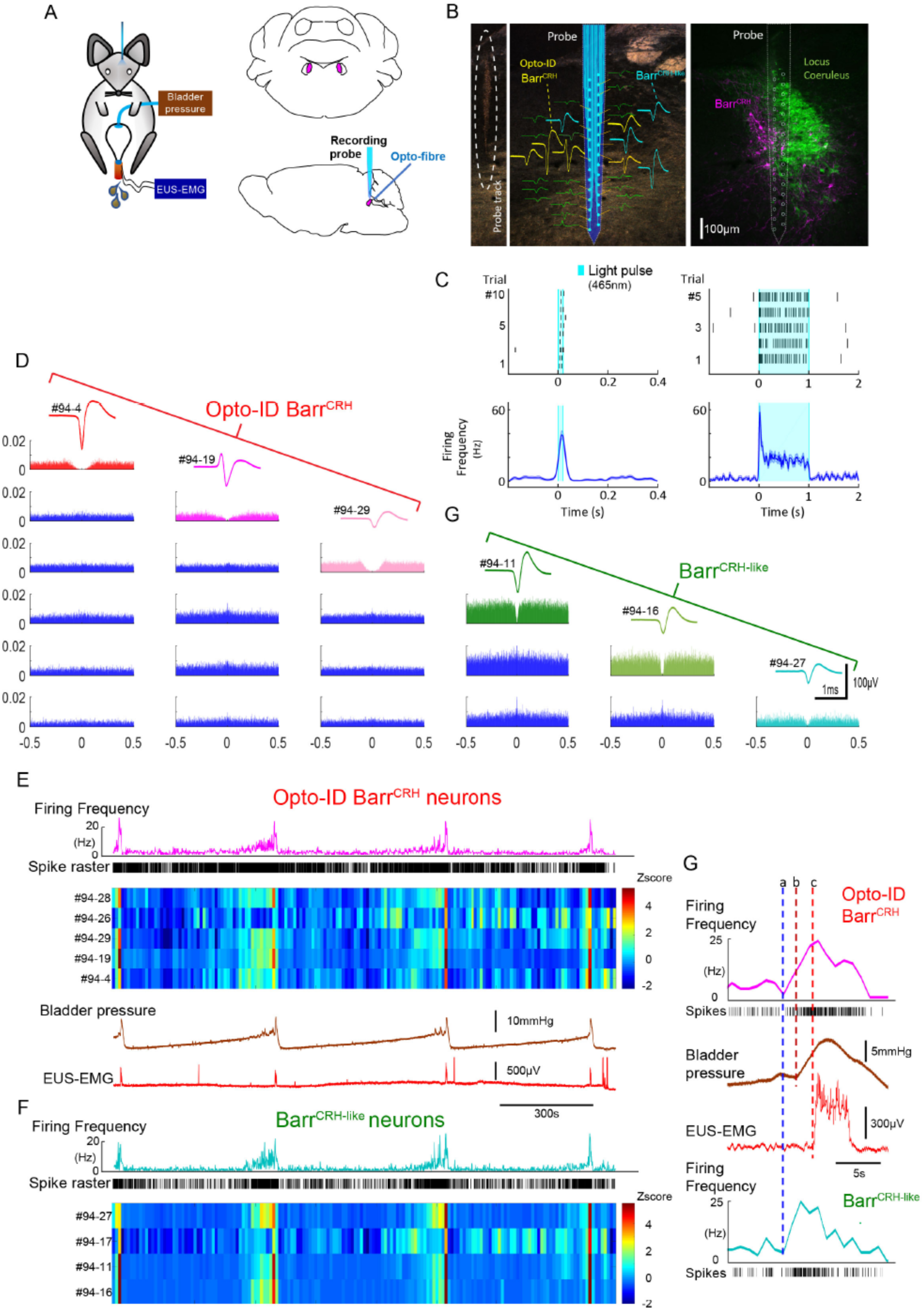
Multiunit recordings of identified Barr^CRH^ neurons. A) Schematic of the experiment with unilateral stimulation and recording of Barrington’s nucleus with simultaneous bladder and EUS monitoring. Note the optical fibre was angled down through the cerebellum to reach Barrington’s whereas the silicon multi-site recording probe was lowered vertically. B) Immunohistochemistry (mCherry - magenta and TH - green) confirming the position of the recording electrode (shown to scale and with its tip at the end of the histological track) and the optical fibre as being either in Barrington’s or the near vicinity (respectively). The spike waveforms of individual units are shown adjacent to their recording site on the probe. Note that the Barr^CRH^ neurons are clustered in probe sites that are located within Barrington’s nucleus. C) Barr^CRH^ neurons were optoidentified by a short latency response to a brief light pulse (20ms) data shown for a single representative unit top left. The population response of identified Barr^CRH^ neurons shown below (*n=12*). Similarly, the response to a 1 second prolonged pulse is shown in the right with the same single unit and the population response from all Barr^CRH^ neurons. Note that they showed an initial high frequency response that decayed to a plateau of ∼20Hz likely reflecting the kinetics of ChR2 currents. D) Auto- and cross-correlations (5ms bin size) of three opto-identified Barr^CRH^ neurons with their average spike waveforms from this recording showing isolation and also a degree of cross correlation at short latency. E) The opto-identified Barr^CRH^ neurons showed a bursting pattern of discharge that was aligned with bladder pressure. The z-scored responses of all Barr^CRH^ neurons in this recording can be seen to have a similar pattern of activity (single representative firing rate plot shown above for reference). F) within the same recording (and from similar probe sites) a further group of neurons was noted (*n=4*) to exhibit a similar pattern of bursting discharge synchronized to the voiding cycle. These cells were termed Barr^CRH-like^ neurons. G) Auto and Cross-correlations of the Barr^CRH-like^ and Barr^CRH^ neurons showed them to have similar properties and evidence of a degree of short latency correlation to other Barr^CRH-like^ neurons and also Barr^CRH^ neurons.

### Spinal CRH inhibits the bladder response to Barr^CRH^ activation

It has been proposed that CRH released from Barrington’s neurons at a spinal level augments bladder pressure responses^39,40^ although others have reported the opposite action^41–43^. If the release of CRH does increase during the micturition cycle, then this might be predicted to act as a positive feedforward mechanism to augment the parasympathetic and hence bladder pressure responses to Barrington’s drive. To test this hypothesis, the effect of intrathecal Astressin (a broad-spectrum CRH antagonist, 5µg in 5µl) on Barr^CRH^ eNVC was assessed through the micturition cycle. Counter to the prediction, Astressin significantly and reversibly increased the amplitude of eNVC, an action that was more pronounced as the bladder filled (333 ± 75 %, *p*=0.008 (*n=7*), 20 mins after Astressin, Figure 4). Intrathecal Astressin also decreased the infused volume required to trigger a void (Figure 4D).

This indicates that CRH is providing a negative feedback signal to limit the extent of the spinal parasympathetic response to Barr^CRH^ neuronal activity (in agreement with^41–43^). Therefore, increased release of CRH cannot account for the augmented responses to Barr^CRH^ activation with progression through the micturition cycle.

### Barr^CRH^ activity anticipates bladder pressure during the micturition cycle

Neural recordings from cats^26^ and rats^30^ indicates that some putative Barrington’s neurons fire intermittently during the storage phase with an increase of firing that occurs around voiding, consistent with a role in mediating the drive to bladder parasympathetic neurons. Recent fibre photometric recordings of Barr^CRH^ neurons, using the genetically encoded calcium indicator GCaMP6, indicate that the activity of these neurons is ‘in phase’ with the micturition cycle^22,23^. However, fibre photometry is unable to resolve the action potential discharge patterns from Barr^CRH^ neurons *in vivo*. As such it has not previously been possible to directly assess the functional relationship between Barr^CRH^ firing and bladder pressure.

Neuronal activity was recorded in the vicinity of Barrington’s using a 32-channel silicon probe to test whether changes in the excitability of Barr^CRH^ neurons during the micturition cycle accounts for the observed variation in the evoked pressure responses of the bladder. An optic fibre was placed above Barrington’s nucleus enabling optogenetic identification (Figure 5A). Recordings were made of cell activity during the normal micturition cycle (with simultaneous bladder pressure and EUS EMG activity) and in response to the application of light stimuli. A total of 113 individual neurons were identified by clustering from recordings made in the vicinity of Barrington’s (*n=3* mice, Figure 5B,C). Definitive opto-identification of Barr^CRH^ neurons (*n=12*) was indicated by reliable short latency spike entrainment to light (20ms pulses, Figure 5C) with time-locked, maintained firing in response to longer light pulses (≥1s, Figure 5C).

These Barr^CRH^ neurons showed a characteristic pattern of activity during the micturition cycle with bursting at the time of voiding (Figure 5E, 20.5 ± 4.1 Hz peak firing frequency). A second population of neurons was recorded with a similar pattern of activity that are henceforth termed Barr^CRH-like^ (*n=32*, Fig 5F, 6A,B) in distinction to the remainder of non-identified neurons (*n=69*). These Barr^CRH-like^ neurons had a short-latency synchrony with the Barr^CRH^ neurons that was evident in cross-correlograms (Figure 5D). Both Barr^CRH^ and Barr^CRH-like^ neurons showed a clear temporal relationship to bladder pressure (Figure 5G) with their firing preceding and ramping up with the pressure during voiding.

**Figure 6.**
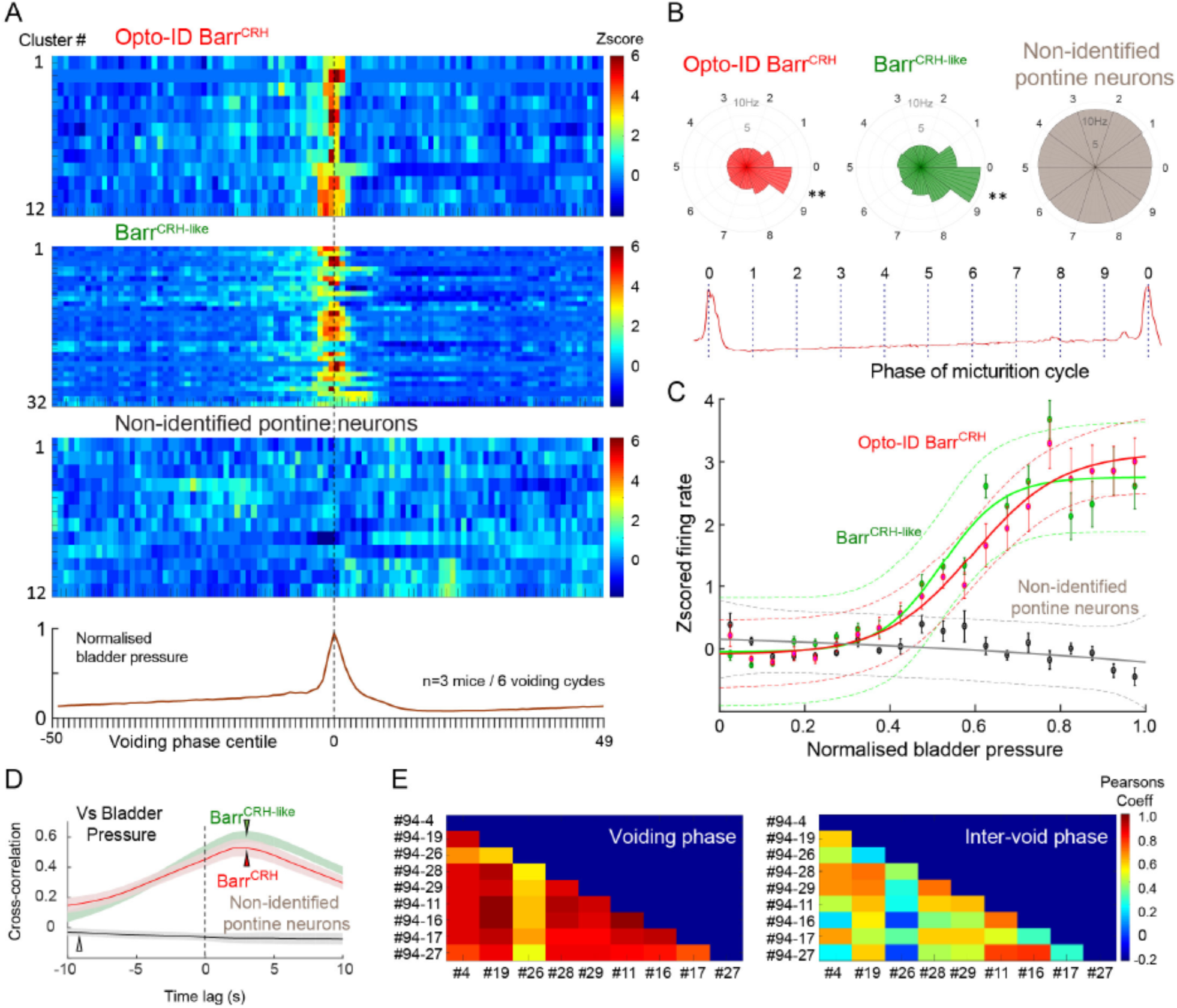
Population dynamics of Barr^CRH^ and Barr^CRH-like^ neurons. A) Silicon probe recordings across mice (*n=3*) with opto-identified Barr^CRH^ neurons (n=12) and Barr^CRH-like^ neurons (*n=32*) showed very similar patterns of firing in relation to the voiding cycle (shown below normalized for pressure and time across 6 cycles). B) Rose plots of firing activity against phase micturition cycle showing that both Barr^CRH^ and Barr^CRH-like^ neurons increase their firing in the phase decile leading up to the void unlike the unidentified neurons. C) Plotting the relationship between firing rate and normalized bladder pressure showed a graded sigmoid relationship with increased firing rate corresponding to higher bladder pressures. No such relationship was seen for the other neurons in the dorsal pons. D) The cross correlation between Barr^CRH^ (and Barr^CRH-like^) neurons and bladder pressure was strongest at a lag of 3 seconds indicating that the bladder pressure follows the change in neuronal firing. E) Colour plots of the Pearson’s cross-correlation coefficient between pairs of the population of Barr^CRH^ and Barr^CRH-like^ neurons is consistently strongest in the voiding phase.

For both the Barr^CRH^ and Barr^CRH-like^ neurons there was a strong sigmoid relationship between bladder pressure and neuronal firing, which was (Figure 6C) not seen in the non-identified group of neurons. The directionality of this influence was investigated by examining the cross-correlation between firing rate and bladder pressure – this indicated that the increases in firing frequency (for both Barr^CRH^ and Barr^CRH-like^ neurons) preceded increases in bladder pressure by ∼3 seconds for both sets of neurons (Figure 6D). These data indicated that the pattern of firing of both Barr^CRH^ and Barr^CRH-like^ neurons anticipated changes in bladder pressure as would be expected for a pre-motor population upstream of bladder parasympathetic neurons. This suggests that the Barr^CRH-like^ neurons are likely to be part of the population of Barr^CRH^ neurons, of which only a subset recorded by the probe are exposed to enough light to be formally opto-identified.

### Barr^CRH^ neuronal excitability is not altered during the micturition cycle

Analysis of spontaneous Barr^CRH^ firing rates over the micturition cycle shows a pattern of activity that is consistent with what would be expected for a high-fidelity controller of bladder pressure. However, this is at odds with our optogenetic activation findings. To resolve this discrepancy the relationship between cycle phase and the light-evoked Barr^CRH^ activity and voiding was examined in more detail.

During all phases of the voiding cycle it was possible to opto-excite Barr^CRH^ neurons and the increase in firing frequency in both absolute and relative terms was independent of the phase of the micturition cycle (ranging from 22.4 ± 7.7 to 24.0 ± 6.5 Hz across micturition phases). These data indicate that the intrinsic excitability of the Barr^CRH^ neurons does not vary across the micturition cycle and that augmentation of the bladder pressure responses (by 17.0±3.9 fold) occurs downstream of the firing output from Barrington’s nucleus.

To further explore this proposition, the relationship between spontaneous NVC (sNVC) and Barr^CRH^ neuronal firing was mapped. A burst of firing in the Barr^CRH^ neurons preceded the sNVC by 2.5-3.0 s – suggesting that they were triggered by a signal from the pons (Figure 7C&D). However, there was only a weak relationship between the magnitude of each Barr^CRH^ burst and the amplitude of the associated sNVC (see Figure 7E). A linear fit of these data indicates that an increase in burst size of 20 spikes (close to the maximum observed range) would only account for 0.5mmHg difference in sNVC size (less than 20% of the observed range of amplitudes). Even this modest relationship was noted to be dependent upon a single outlier value of a large NVC occurring close to a void (circled). Again, this finding is consistent with Barr^CRH^ providing a trigger signal rather than a pre-motor drive which determines the amplitude of the bladder contraction.

**Figure 7.**
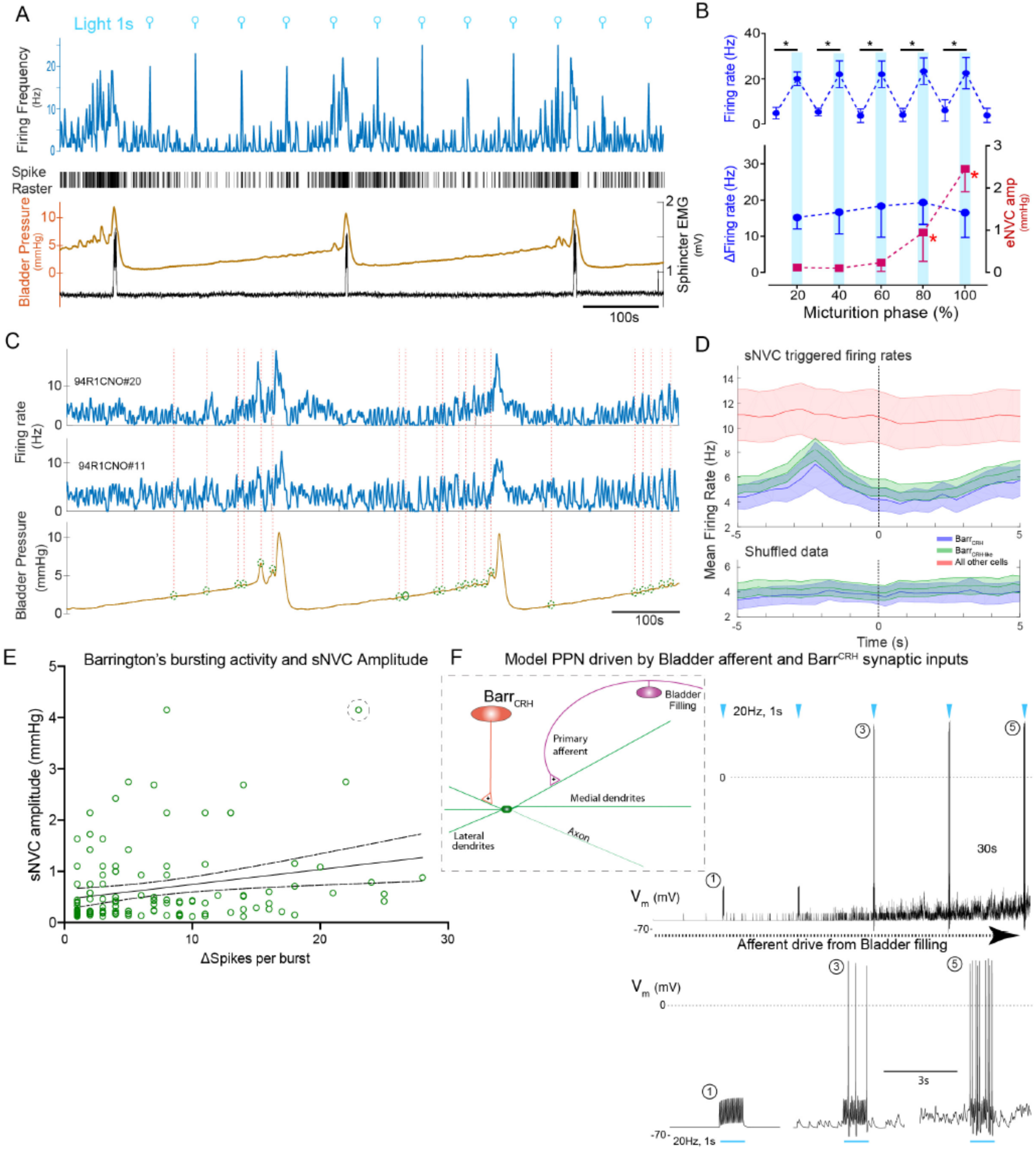
Barr^CRH^ neuronal activity conditionally drives bladder pressure. A) Optogenetic stimulation of Barr^CRH^ neuron showing the transient increases in firing evoked by light pulses (1s x 465nm) applied at different points of the micturition cycle. B) Pooled data from all Barr^CRH^ neurons (*n=12* across 3 mice) showing that there was no difference firing (either the peak firing rate (upper) or the change in firing (lower)) evoked by light across the phases of the micturition cycle. In contrast the amplitude of the eNVC (see figure 3) increases markedly across micturition cycle. C) Spontaneous NVCs were identified using a peak find algorithm (amplitude 0.1-4mmHg) and were noted to be preceded by a burst of Barr^CRH^ activity. D) Barr^CRH^ and Barr^CRH-like^ neurons showed a burst of firing between 1.5-3s before the onset of sNVCs (unlike the unidentified population and also not seen in the shuffled data). Mean firing rates ± S.D, 0.5s bins. E) The number of spikes in each Barr^CRH^ burst only showed a weak correlation (slope 0.03mmHg/spike) with the amplitude of the following sNVC. This weak relationship was lost if a single outlier point was excluded (ringed). F) Model of a parasympathetic preganglionic neuron (implemented in NEURON) with a synaptic drive from Barrington’s nucleus. To mimic the change in excitability through the micturition cycle an incrementing depolarising current was added. This shows that a 20Hz stimulation of Barr^CRH^ (mimicking opto-activation) evoked no parasympathetic output at the start of the cycle but this increases to 8-10hz by the end of the cycle.

### A spinal gate for the Barr^CRH^ drive is opened by bladder distention

This indicates a model of autonomous micturition where a spinal circuit gates the output to the bladder (shown schematically in Figure 7F). The Barr^CRH^ – parasympathetic - bladder afferent component of this circuit was modelled in NEURON using an existing preganglionic neuronal model^44^ and a combination of a fast, excitatory synaptic drive descending from Barrington’s plus a bladder afferent synaptic drive (based on recordings of pelvic nerve afferents from Ito *et al*^34^). The incrementing frequency of afferent drive, as the bladder fills, leads to summation and a maintained membrane depolarisation that increases parasympathetic excitability. The resulting output from the parasympathetic neuron when driven by Barr^CRH^ (with a mimicked 20Hz optogenetic drive) was strongly dependent upon the phase of the micturition cycle with a ∼10-fold increase over the voiding cycle which closely parallels the experimental data. Note also that in the early phase of the voiding cycle the Barr^CRH^ input is unable to evoke action potentials – thus producing ‘failures’.

### An inferential model of autonomous micturition

The observations described suggest a new integrated model of the micturition cycle which incorporates the observed drive from Barr^CRH^ neurons and the known afferent feedback from the bladder (Figure 8A and methods). This afferent feedback governs both the excitability of the spinal parasympathetic neurons (demonstrated in the NEURON model above) and the output of a synaptic generator driving Barr^CRH^ activity. The resulting feedback loop (depicted schematically in Figure 8B) closely reproduces characteristic of the observed micturition cycle with graded NVCs, periodic voids and patterns of Barr^CRH^ firing.

**Figure 8.**
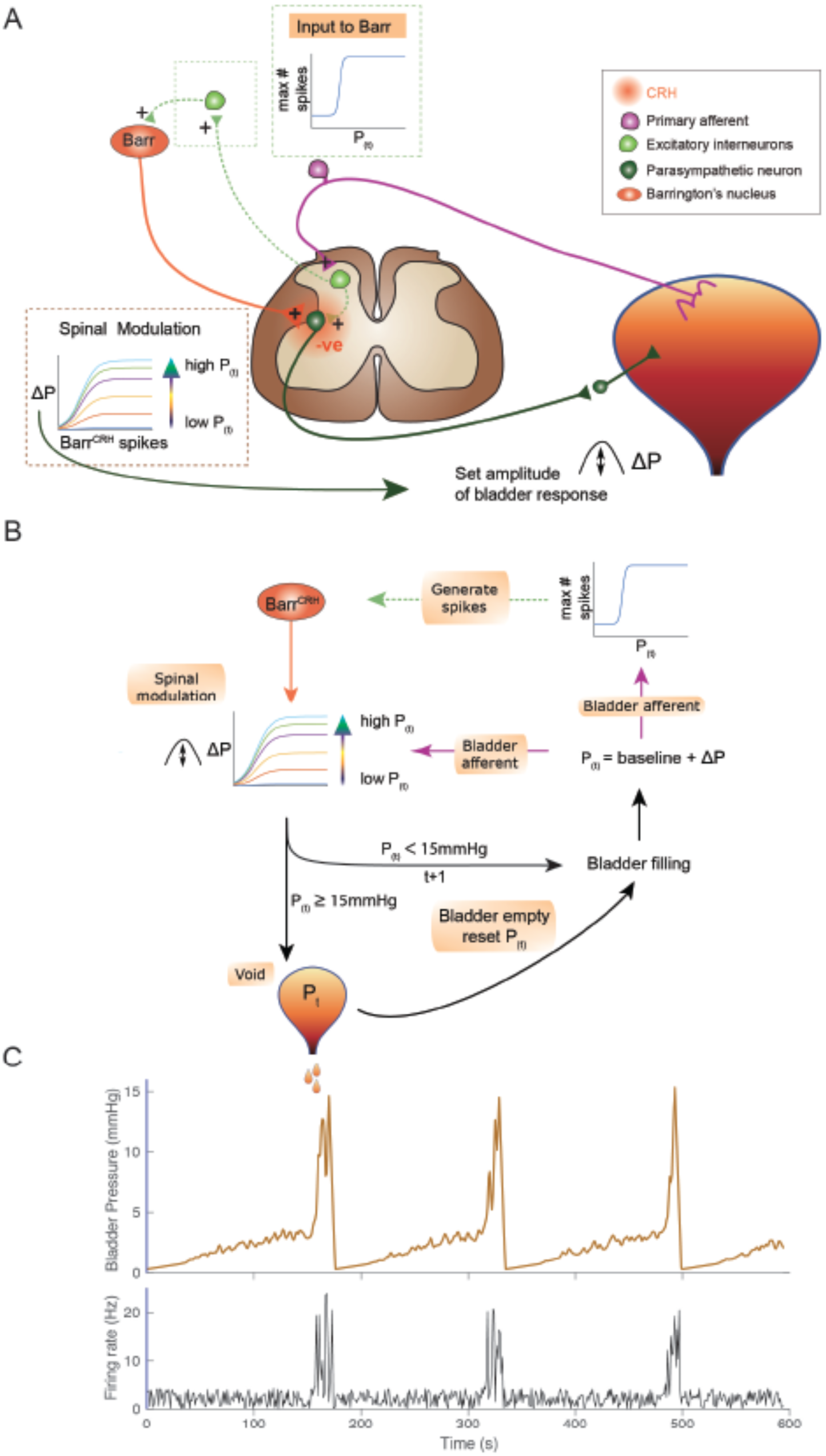
An inferential model of autonomous micturition. A) Schematic of the descending input from Barrington’s nucleus to the bladder parasympathetic neurons. Also showing the excitatory input to parasympathetic neurons of the bladder afferent information – shown as mediated via a segmental excitatory interneuron. Note that the Barr^CRH^ neuron has both a fast, excitatory transmitter (presumed glutamate) as well as an inhibitory action mediated by released CRH – possibly acting via local inhibitory interneurons (not shown). The inset boxes show the logistic relationships linking activity of Barr^CRH^ neurons and spinal excitability to the current bladder pressure (from model in B). B) Flow chart showing processing steps in inference model of micturition. C) Output from the model showing incrementing bladder pressure with NVCs over 3 micturition cycles with the associated Barr^CRH^ firing that generates the NVCs and the voids.

The varying excitability of spinal parasympathetic neurons is represented in the integrated model by a pressure-modulated logistic relationship which determines the change in bladder pressure generated from a given level of Barr^CRH^ firing. The afferent drive also determines the probability of a high frequency Barr^CRH^ discharge in a given epoch, again via a logistic relationship – modelling a pressure dependent synaptic drive for Barr^CRH^. This synaptic generator is commonly believed to be within the PAG^2,5,45^; however, recent studies indicate it could also be located in the cortex or hypothalamus^22,24^.

This circuit organisation generates NVCs : dynamic perturbations whose magnitude and frequency increase with progress through the micturition cycle. As pressure increases these contractions become more frequent and higher in amplitude – eventually summating to cause sustained increases in bladder pressure. This in turn increases the rate of firing of Barr^CRH^, making further contractions more likely, and shifting the system into a positive feedback loop in which pressure rapidly increases. A void occurs when the pressure threshold is exceeded (in the model, this is set at 15mmHg) which is presumed to be effected via a spinal mechanism.

In line with experimental data, attenuation of the variance in Barr^CRH^ firing (underpinning the NVCs) delays the time to void – indicating their importance in the process (Supplementary Fig 6A). Similarly, augmenting the spinal parasympathetic sensitivity to the Barr^CRH^ drive (as seen experimentally with intrathecal Astressin) increases the amplitude of the NVCs and shortens the inter-void interval (Supplemental Fig 6C). We note that additional drive into the Barr^CRH^ neurons as is proposed to come from higher centres with voluntary voiding would increase the variance and could trigger voiding earlier. This effect is demonstrated with the simulated optogenetic drive of Barr^CRH^ neurons (20Hz x 1s, Supplemental Fig 6B) which produces both failures, eNVCs and triggers voids earlier in the cycle than would otherwise have happened.

### Discussion

These findings indicate that Barr^CRH^ neurons do play a critical role in micturition. However, the activity of Barr^CRH^ neurons, is not a simple switch mechanism for voiding nor do they provide a direct drive to bladder pressure (as might be expected for an autonomic pre-motor command neuron). Instead these neurons play a more nuanced, probabilistic role. Their influence on the bladder depends on the state of priming of the downstream parasympathetic motor circuit. This identifies the Barr^CRH^ neurons as being the efferent limb of an inference circuit that assays bladder state repeatedly during the storage phase of the cycle. When the threshold for voiding is reached, they generate a high-fidelity motor signal through a regenerative feedback loop that drives the bladder contraction required for voiding.

This operating principle fits with a modular hierarchical hypothesis for the organisation of the micturition circuit^2,5,32^ with a primary spinal circuit providing a basic functionality, evident in the neonatal rodent^46–48^ and indeed other mammals including humans, that has little context-sensitive control. The timing of micturition in immature rodents is often triggered by maternal stimulation of the perineum (although interestingly this is unsuccessful if applied when the bladder is <50% full^48^). With development, the spinal micturition mechanism is believed to fall progressively under the descending control of Barrington’s nucleus (both for voluntary and autonomous voiding). We suggest that such descending control provides an internalised signal, replacing the need for additional external peripheral sensory input, to trigger the void. Dysfunction of this descending control system, as is seen following spinal cord injury, results in a loss of voluntary control and initially in a complete loss of continence, but this tends to be restored as the spinal micturition reflex re-emerges (albeit in a poorly co-ordinated manner). This situation was mimicked experimentally herein by the chemogenetic inhibition of Barr^CRH^ neurons – leading to a prolongation of the inter-void interval and retained volumes with progressive bladder distension – indicating that this is a necessary and critical component of the micturition circuit.

### Parallel circuits within Barrington’s nucleus regulating urination

We used cystometry in anaesthetised mice to examine the role of Barr^CRH^ neurons specifically in the core processes of autonomous micturition in the absence of behavioural influence. Rodents also use social urine scent marking, for example male mice use strategic urine “spotting” to express their dominance and territorial ownership^49,50^. Although autonomous micturition and scent marking with urine are related processes, likely with some shared physiology, there are also differences in the patterns of urination with greater frequency (>10 urine spots per minute^23^) and accordingly smaller volumes in each urine spot compared to a primary void (typically 80-120ul)^9,51–53^. A role has recently been described for Barr^ESR1^ neurons in social urine spotting evoked by female urine^23^. These Barr^ESR1^ neurons preferentially target a spinal inter-neuronal circuit that is proposed to be involved in generating the bursting drive to the EUS as well as causing a bladder contraction. In contrast, in the same study, the Barr^CRH^ neurons were reported to be relatively ineffective in generating voids under similar conditions (without active filling of the bladder)^23^. However, our study shows that Barr^CRH^ neurons can effectively generate co-ordinated voiding – with the characteristic pattern of external urethral sphincter bursting – but only when the downstream spinal circuit is primed i.e. with a nearly full bladder. This indicates that the drive from Barr^CRH^ needs to be integrated at a spinal level with feedback from the bladder afferents in order to generate a complete void. When the bladder is partially filled, then activation of the Barr^CRH^ neurons evokes bladder contractions (eNVC) without any sphincter activity, suggesting that the drive to the sphincter is also dependent on the state of a downstream pattern generator and is not directly engaged by the firing of Barr^CRH^ neurons at all stages of the micturition cycle. It is also noteworthy that there was no evidence for the involvement of Barr^CRH^ neurons in the control of the distal colon, suggesting they are bladder specific.

Given the mechanistic differences between the processes of autonomous micturition and voluntary scent marking in males, it is quite likely that there are distinct circuit drives for each type of urination. This would be consistent with the proposition that there are parallel pathways from Barrington’s to the downstream spinal pattern generators: the Barr^ESR1^ neurons driving spotting behaviour which can be triggered irrespective of the degree of fullness of the male mouse bladder; and the other mediated by Barr^CRH^ neurons which requires the bladder to be distended before a void can be generated. This may also explain why chemogenetic inhibition of the Barr^ESR1^ but not Barr^CRH^ neurons blocked spotting whereas similar inhibition of Barr^CRH^ neurons inhibited autonomous micturition in the current study.

### Barr^CRH^ neurons provide a probabilistic, spinally-gated drive to bladder pressure

The pattern of firing activity seen in optogenetically-identified Barr^CRH^ neurons is similar to that previously noted in recordings from Barrington’s nucleus in the conscious rat^30^ and is also reminiscent of a subset of the neurons identified in anaesthetised or decerebrate rats and cats that showed a ramping activity with voiding^25–29^. The bursting activity seen in the Barr^CRH^ recordings clearly precedes the changes in bladder pressure (and with a similar lag to bladder pressure response to that found from optogenetic activation of Barr^CRH^ neurons) indicating that they are driving rather than responding to the changes in pressure. The Barr^CRH^ neurons showed a pattern of spiking activity that is also consistent with that noted from the Ca^2+^ imaging recordings seen with fibre photometry in mice^22,23^ although we can see both the temporal precedence and that this activity decays promptly at the end of the void. It is also worth noting that in none of these recordings (from Barr^CRH^ or indeed any of the neurons in the vicinity) was any pattern of activity seen that resembled the high frequency bursting of the urethral sphincter seen in mice and rats – to date such activity has never been observed in any of the recordings from Barrington’s which is consistent with the idea that it is generated from a spinal motor pattern generator such as the LSCC^38^.

The probabilistic nature of the influence of Barr^CRH^ neurons on bladder pressure seems initially at odds with the clear relationship between their activity and bladder pressure noted herein (and previously in the E2 class of neurons recorded in rat Barrington’s^28^). However, this relationship only holds in the late stage of the micturition cycle when Barr^CRH^ neurons do indeed act as a tightly-coupled, direct command neuron. This is not the case during the early phases of the micturition cycle when there is a weak relationship between the activity of Barr^CRH^ neurons and the bladder pressure, in the extreme case leading to “failures” of stimulation to evoke any contraction. Our recordings indicate that this happens downstream of Barr^CRH^ at a spinal level, as the ability to optogenetically drive Barr^CRH^ is unchanged and spinal activation of Barr^CRH^ axons also shows failures. We propose a model for this action that integrates an incrementing and summating, but still sub-threshold, afferent drive from the bladder to PPN that enables a phasic burst of activity from Barr^CRH^ to generate progressively larger numbers of action potentials and hence contraction when the bladder is sufficiently filled. In support of this idea, previous electrical stimulation studies of Barrington’s nucleus in the rat indicated that the degree of excitation of the parasympathetic motor outflow to the bladder was strongly dependent upon the degree of bladder filling^54^. The mechanisms enabling such priming of the parasympathetic control circuit will merit further investigation at a spinal level.

### The role of spinal CRH in the regulation of micturition

The inhibitory action of CRH released from Barr^CRH^ neurons on micturition at a spinal level initially appears counterintuitive given the overall excitatory effect of Barr^CRH^ neurons (mediated via fast glutamatergic signalling,^22^) on micturition and the known excitatory effects at a cellular level of CRH receptor activation^55^. However, a similar inhibitory spinal action of CRH on micturition has been reported^41–43^. This inhibition may act to suppress the segmental excitatory activity in the spinal parasympathetic circuit. This process may also be involved in the transition from the immature spinal voiding circuit in the neonate that is supplanted by the top down requirement for Barrington’s signals. The mismatch between the differential time-course of action of CRH-mediated inhibition (metabotropic) and the fast, glutamatergic excitation (ionotropic) may enable the initial rapid excitation of parasympathetic preganglionic neurones at the onset of voiding, but may also in turn act to help terminate voids and facilitate the unopposed relaxation of the bladder. The spinal mechanism of CRH actions at a spinal circuit level merits further investigation as this constitutes an intriguing target for therapeutic intervention allowing modification of the gain of the micturition reflex in disease states.

### A novel inferential model of autonomous micturition

We noted that bursts of activity in Barr^CRH^ neurons precede both voids and also NVCs. There has been considerable debate about the origin of NVCs with respect to whether they are intrinsically generated by the bladder and their functional significance although there is a suggestion that they provide a means to infer the degree of bladder filling^56^. An increase in their frequency and amplitude has been linked to diseases of the LUT^57^ and also with loss of descending control from the brainstem ^6^ which could conceivably be related to the loss of CRH-mediated spinal inhibition. There is also evidence of their peripheral generation by the bladder early in development which becomes less coordinated in adult bladder ^58^. We provide evidence that NVCs are generated by the “noisy” probabilistic drive from Barrington’s that repeatedly assesses the status of the spinal circuit during each micturition cycle and that the magnitude of the bladder pressure response reflects the phase of the micturition cycle. The resulting afferent signal provides an active way of inferring the degree of bladder fullness (analogous to the ‘sampling’ that assesses rectal fullness ^59^) and could prime the neural control circuits and indeed could conceivably provide a stream of information that may enable a conscious awareness of bladder fullness and the ability to make volitional predictions about the need to void. We also note the homology with the development of other motor systems where spontaneous motor activity, initially generated in the periphery, becomes progressively embedded centrally as motor representations in the nervous system with developmental maturation^60^.

Our model of such a circuit organisation with afferent feedback from the bladder both priming the spinal parasympathetic motor circuit and also determining the magnitude of the drive from Barr^CRH^ neurons (perhaps via integration by the PAG which functions as a probabilistic firing switch) recapitulates many of the observed features of autonomous micturition. The generation of NVCs provides inference about the degree of bladder fullness and the afferent signal advances the progression through the cycle. The spinal priming mechanism enables a regenerative burst of activity from Barr^CRH^ to drive the voiding contraction and the modelled release of spinal CRH that follows such a large discharge serves to reset the spinal circuit enabling passive filling to resume. A feature of this circuit organisation is that a direct volitional drive to Barr^CRH^ from cortex as recently reported^24^ would not be subject to the probabilistic firing switch at a supraspinal level and could therefore trigger a void earlier in the cycle if behaviourally appropriate albeit still contingent on the priming status of the spinal gate. Hypothetically, a parallel synaptic drive from the Barr^ESR1^ neurons^23^ that was stronger than the Barr^CRH^ neurons could also generate parasympathetic activity without requiring co-incident afferent activity – hence bypassing the gate to produce urine “spotting” on behavioural demand.

On this basis we conclude that the Barr^CRH^ neurons form a key component of the micturition circuit that generate a pre-motor drive to the bladder late in the cycle. The recording and stimulation data suggest that this drive is not generated by a burst generator residing within this cell population but is a product of the integration of inputs from both bladder sensory afferents and upstream centres such as the PAG but also including hypothalamus and motor cortex ^22,24^. We predict that failures of control at this key integrating locus are likely to be involved in both acute disorders of lower urinary tract function such as retention as well as in chronic diseases like nocturnal enuresis and overactive bladder syndrome where there is dissociation of detrusor contractions and sphincter relaxation.

## Acknowledgements

Funded by US NIH R01 DK098361 (Anthony J. Kanai, Marcus J. Drake, Christopher H. Fry, Anthony E. Pickering). ACS is supported by the Wellcome Trust PhD programme in Neural Dynamics (ref. 108899/Z/15).

## Online methods

### Experimental model and subject details

#### Mice

All experiments and procedures conformed to the UK Animals (Scientific Procedures) Act 1986 and were approved by the University of Bristol Animal Welfare and Ethical review body. Mice were group housed, with food and water available *ad libitum* and on a 12 hr/12 hr light/dark cycle.

Gene expression was restricted to Barr^CRH^ neurons using knock-in mice (of both sexes aged 3-8 months old) with an internal ribosome entry site (ires)-linked Cre-recombinase gene downstream of the CRH locus (CRH^Cre^ mice^1,2^, Jax Laboratory #012704).

### Methods

#### Viral vectors

The serotype 2 recombinant AAV-EF1*α*-DIO-hChR2(H134R)-mCherry^3,4^ (1.6 x 10^12^ viral genomes / ml) was obtained from University of North Carolina vector core facilities (a gift from Karl Deisseroth), whereas serotype 2 AAV-hSyn-DIO-hM4Di-mCherry^5^ (7 x 10^1^^2^ vg / ml) was obtained from Addgene (a gift from Brian Roth).

#### Stereotaxic Intracranial Injections to Barrington’s nucleus

To target Barr^CRH^ neurons, homozygous CRH^cre^ mice were anaesthetised with ketamine (70 µg/g) and medetomidine (0.5 µg/g) and placed in a small animal stereotaxic frame (Kopf, USA) with a drill-injection robot attachment (Neurostar, Germany). After exposing the skull under aseptic conditions, a small burr hole was drilled and AAVs were injected (600nl total volume per side) unilaterally or bilaterally through a pulled glass pipette at a rate of 100nl/ml. Injection coordinates for Barrington’s nucleus were 5.3 mm posterior to bregma, 0.70 mm lateral and 3.5 mm below brain surface. After surgical procedures, all mice were returned to their home cage for at least 21 days for recovery to maximise protein expression.

#### Optogenetic Activation

Adult CRH^Cre^ mice had AAV-DIO-ChR2-mCherry (1.6 x 10^12^ vg/ml) or AAV-DIO-hM4Di-mCherry (as control) injected into Barrington’s and they were used in experiments at least 3 weeks after vector injections. Light from a 465 nm LED (Plexon, Dallas USA) was delivered in pulses with a maximum duty cycle of 50%. The light train was delivered once every 60 s for fixed-interval stimulation, or at randomised intervals between 30 s and 90 s. The light power exiting the fibre tip was set at approximately 10 mW and was measured before and after each experiment. For bilateral simultaneous opto-activation a dual fibre implant was used (Doric, DFC 200/250-0.66 15mm DF1.4 C60) and coupled via a dual fiber optic cable to two separate LEDs.

For light delivery to the spinal cord, soft tissue was removed between T11 and T12 vertebral spines after skin incision. The exposed spinal cord was illuminated using a 473nM laser (PhoxX, Omicron, Germany) via a bare ended fibre (Thorlabs, 400µm) positioned above the cord and delivered in 20ms pulses at 20Hz for 5s or a prolonged pulse of 1000ms. The light power at the fibre tip was 29±0.3mW.

#### Chemogenetic Inhibition

To inhibit Barr^CRH^ neurons, CRH^cre^ mice were bilaterally injected with AAV-DIO-hM4Di-mCherry (7.0×10^10^ vg/ml) into Barrington’s nucleus (as described above) and allowed at least 3 weeks of recovery (control mice had AAV-DIO-ChR2-mCherry injected). Cystometry was performed as described below and intraperitoneal CNO (5mg/kg, 1mg/ml stock) or saline (as control) was applied after obtaining > five baseline micturition cycles. In an initial set of experiments, saline was continuously infused to the bladder around the time of CNO injection to investigate the effects on mice micturition. Subsequently, to determine the CNO effect on threshold for micturition, a cyclical infuse and hold protocol was adopted whereby saline infusion was stopped at the threshold for voiding and then held at that volume for 10 minutes or until a void occurred before emptying the bladder and restarting the infusion phase.

#### Cystometry, Electromyography and distal colonic manometry.

Mice were anesthetised with urethane (0.8-1.2mg/kg) and the bladder was exposed via a 2 cm midline abdominal incision. A flanged catheter (PE50) was secured with a purse-string suture into the bladder and connected to a syringe pump and pressure transducer. The infusion rate was adjusted on an individual mouse basis (10-40 µl/min) to produce an equivalent proportionate speed of fill to threshold for voiding (typically 600 sec) taking account of differing bladder volumes of the mice. External urethral sphincter (EUS) was recorded with insulated stainless steel wires, bared at the tip (0.075mm, AISI316 Advent) inserted through a 30-G needle bilaterally into the EUS just proximal to the pubic symphysis. A balloon catheter (2.5mm diameter x 12mm when fully distended, Medtronic Sprinter) was inserted into distal colon and the tip of balloon was placed 40mm from the anus. To monitor colonic pressure the balloon catheter was filled with distilled water.

More than 1 hour after starting saline infusion into the bladder and once a regular rhythm of micturition cycles was established. The following variables were measured (averaged over at least three voiding cycles):

- Basal pressure was taken as the lowest bladder pressure reached after a void.
- Voiding threshold was the bladder pressure when the EUS-EMG started bursting, indicating the initiation of voiding.
- Micturition pressure was the peak bladder pressure achieved during voiding (bursting phase of the EUS-EMG).
- Non-voiding contractions (NVCs) were identified as discrete increases in bladder pressure (>1 mmHg) observed during the filling phase in voiding preparations.
- Bladder compliance was defined as bladder capacity / (threshold - basal pressure) (µl/mmHg) during filling.

#### Pithed, Decerebrate Arterially-Perfused mouse preparation

The pithed DAPM preparation was used to examine the influence of bladder filling on pelvic nerve stimulation-evoked bladder contractions. The methods were as previously described^6,7^ but in brief, mice were terminally anaesthetised with isoflurane, disembowelled through a laparotomy and the bladder was cannulated. The mouse was then cooled, exsanguinated, decerebrated and its spinal column was pithed to remove all central neuronal control. It was then moved to a recording chamber, perfused through the heart with warm (32°C) Ringer’s solution (composition (mM): NaCl (125), NaHCO_3_ (24), KCl (3.0), CaCl_2_ (2.5), MgSO_4_ (1.25), KH_2_PO_4_ (1.25); glucose (10); pH 7.35–7.4 with 95% O2/5% CO2). Ficoll-70 (1.25%) was added as an oncotic agent to the perfusate. The flow rate was adjusted (from 15 to 20 ml/min) to achieve a perfusion pressure of 50-60mmHg. The pelvic nerve was identified, traced proximally and cut allowing the distal end to be aspirated into a bipolar stimulating electrode. Stimuli (10V, 1ms, 4-10Hz for 3 seconds) were applied to the nerve. The bladder was filled with saline to perform cystometry as above with filling limited to a ceiling pressure of 15mmHg. The effect of pelvic stimulation on bladder pressure was examined with different degrees of bladder filling (0-70µl).

#### Extracellular recordings and signal acquisition

Recordings were made from the Barrington’s nucleus using a 15 μm thick silicone probe with 32 channels (NeuroNexus, Model: A1×32-Poly3-10mm-25 s-177-A32). Each channel (177µm^2^) was spaced from the neighbouring channels by 50 μm. A reference electrode (Ag/AgCl) was inserted into the scalp. The probes were connected to an amplifier-digitising headstage (INTAN, RHD2132). The signals were amplified and filtered (100Hz-3 kHz) and digitized at 30kHz before being processed and visualised online within the Open Ephys system.

#### Anatomical tracing studies

To investigate the Barr^CRH^ projection to the spinal cord, Cre-dependent AAV (AAV-EF1*α*-DIO-ChR2-mcherry) was unilaterally injected to Barrington’s nucleus in CRH^CRE^ mice. After a minimum of four weeks the mice were killed, and perfusion fixed for immunohistochemistry. To examine the Barr^CRH^ projection into spinal cord, 40µm transverse sections were taken from T11 to S2, processed for mCherry and Choline acetyltransferase immuno-(to demarcate motoneurons) followed by confocal imaging (detailed below).

#### Immunohistochemistry of brain and spinal cord

Mice were killed with an overdose of pentobarbital (20 mg per mouse, i.p; Euthetal, Merial Animal Health) and perfused trans-cardially with 4% formaldehyde (Sigma) in phosphate buffer (PB; pH 7.4, 1 ml/g). The brain and spinal cord were removed and post-fixed overnight before cryoprotection in 30% sucrose in phosphate buffer. Coronal tissue sections were cut at 40 µm intervals using a freezing microtome and left free floating for fluorescence immunohistochemistry. Tissue sections were blocked and incubated in phosphate buffer containing 0.3% Triton X-100 (Sigma) and 5% normal donkey serum (Sigma). Incubated on a shaking platform with primary antibodies for 14–18 hr at room temperature. After washing, sections were then incubated for 3 hr with appropriate Alexa Fluor secondary antibodies.

A Leica DMI6000 inverted epifluorescence microscope equipped with Leica DFC365FX monochrome digital camera and Leica LAS-X acquisition software was used for widefield microscopy. For confocal and tile scan confocal imaging, a Leica SP5-II confocal laser-scanning microscope with multi-position scanning stage (Mӓrzhӓuser, Germany) was utilized. Primary antibodies used were rabbit anti-mCherry (1:4,000; Biovision), sheep anti-tyrosine-hydroxylase (1:1,000; AB1542, Millipore) and goat anti-ChAT (1:250; AB144P, Millipore). Alexa Fluor 488-conjugated donkey secondary antibodies were used against goat IgG (1:500; Jackson ImmunoResearch) and sheep IgG (1:400; Jackson ImmunoResearch). Alexa Fluor 594-conjugated donkey secondary antibody was used against rabbit IgG (1:1000; Invitrogen).

#### Model design

A model of the integration of the synaptic drive to the bladder parasympathetic preganglionic neurons in the spinal cord was constructed using NEURON^8^. The preganglionic neuron was based on using an existing preganglionic neuronal model^9^ that was modified slightly to have a resting potential of ∼-65mV by altering the leak conductance reversal potential. A synaptic drive from Barr^CRH^ neurons was modelled by adding an EXPSYN to the soma which was driven with trains of action potentials (20Hz x 1s) generated from a NETSTIM to mimic an optogenetic stimulus of Barr^CRH^. The synapse was subthreshold if triggered alone as an input despite a modest degree of summation (∼20%) when driven at 20Hz. A second fast excitatory synaptic drive to model a bladder afferent was added as an EXPSYN to a proximal medial dendrite. This was driven with an incrementing frequency of action potentials (from a NETSTIM based on recordings of pelvic nerve afferents from Ito *et al*^7^) to model bladder distension-evoked increase in afferent firing over a period of 5 minutes to represent a typical micturition cycle. The model files are available on ModelDB (TBA).

The model of autonomous micturition employs an iterative algorithm in which bladder pressure and Barr^CRH^ firing rate are updated in each cycle (assumed to last 1s). At each time point, bladder pressure is incremented by 0.015mmHg (modelling a continuous infusion). The level of pressure in the bladder is used to determine the maximum level of Barr^CRH^ firing via a logistic function (modelling an afferent input):

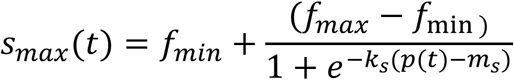

Where *s_max_* is the maximum spiking rate, and *f_max_* and *f_min_* represent the maximum and minimum levels of firing – here set to 25 and 4.3Hz respectively, based on spiking rates seen in unit data recordings. The gradient and mean were set to *k_s_* = 4 and *m_s_* = 4.5. Actual spiking rates were then generated probabilistically by sampling from a random uniform distribution with a maximum level set by *s_max_*.

The change in bladder pressure produced by the firing of Barr^CRH^ neurons was determined by a logistic function modulated by bladder pressure (representing the level of parasympathetic neuron excitability at the level of the spinal cord):

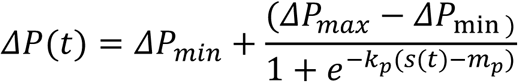

Where *s* is the Barr^CRH^ firing rate and *ΔP_max_, ΔP_min_* are the maximum and minimum amplitude of bladder contractions, and *K_7_* = 0.5 and *m_7_* = 6. The maximum change in pressure depends on the current bladder pressure such that:

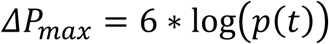

The output of these equations is shown in Figure 8 and Supplementary Figure 6. Bladder contractions are modelled with a Gaussian bump function, of amplitude *ΔP_max_* and variance 2s. These had a of duration 6 timepoints (approximating the characteristics of non-voiding contractions observed experimentally). At the beginning of each cycle, the current pressure is calculated by summing the baseline pressure (including incrementing due to constant filling) with increases in pressure caused by Barr^CRH^ triggered bladder contractions generated in the current and past cycles.

Voiding is triggered at a threshold pressure of 15mmHg, at which point bladder pressure is decremented at 2mmHg per cycle (as the bladder empties) until a baseline pressure of <0.3mmHg is reached and the cycle restarts. The model is coded in MATLAB and has been deposited in the GitHub repository.

In cycles where optogenetic activation of Barr^CRH^ firing was simulated, the firing rate *s* was set to 20Hz. To simulate intrathecal Astressin, the mean of the logistic curve for *ΔP(t)* was reduced to *m*_p_ = 5 To simulate attenuation of NVCs, the variance in the level of Barr^CRH^ firing was reduced by 80% but without any change in the mean level of firing.

### Data analysis

#### Clustering of multiunit data, waveforms, auto / cross correlations and calculation of firing rates

Multiunit data was recorded on a 32-channel silicon probe (NeuroNexus, Model: A1×32-Poly3-10mm-25 s-177-A32), and clustered using spike sorting framework ‘Kilosort’ ^10^. The sorting analysis was carried out using the facilities of the Advanced Computing Research Centre, University of Bristol - http://www.bristol.ac.uk/acrc/. Manual curation of clusters was performed in ‘Phy’ (https://github.com/kwikteam/phy-contrib) in order to select only well isolated units with clear refractory periods, and to remove artefacts. All further analysis of spike trains and cluster characteristics was carried out in MATLAB.

The centre channel of each cluster was defined as the probe channel on which the waveform was recorded with the maximum range. Representative waveforms were extracted on the centre channel for each cluster by sampling 2000 spikes from the group (if the cluster had fewer than 2000 spikes present, all spikes were used (Figure 5B, D). Autocorrelations and cross-correlations were calculated by binning spike trains (1ms bins) and using the MATLAB function ‘xcorr’ (Figure 5D). Smooth firing rates during laser stimulation events were calculated by convolving the spike train with a normalised Gaussian of standard deviation of 30ms, using the MATLAB function ‘conv2’. (Figure 5C). Where z-scored firing rates were required, the MATLAB function ‘zscore’ was used in with binned spike counts.

#### Analysis of bladder pressure and spike trains

Bladder pressure and external urethral sphincter EMG were recorded in Spike2 (CED) and analysed in MATLAB. To compare changes in bladder pressure over multiple recordings, bladder pressure was normalised to between 0 and 1 in each recording, where 1 represents the maximum pressure recorded during the experiment (Figure 6A). The times of voids were identified using the MATLAB function ‘findpeaks’; correct identification was verified by eye. Where the bladder pressure was split into phases of the voiding cycle, the period between successive voids was split into the required number of equal time intervals (100 phases in Figure 6A, 10 phases in Figure 6B). Spike counts during these phases were converted to firing rates by dividing by the width of the relevant time window. In Figure 6A these were also z-scored to enable comparison between multiple voiding cycles recorded in different animals.

Where voiding / inter-voiding periods were used (Figure 6E), ‘voiding periods’ were defined as a window of 15 seconds either side of the peak of bladder pressure during a void; ‘intervoid periods’ consisted of all remaining times.

Cross correlograms and Pearson correlation coefficients between spike count and bladder pressure were calculated by downsampling both data to a sampling rate of 1hz (i.e. 1s bins for the spike count, 1hz sampling for the bladder pressure), z-scoring and using the MATLAB function ‘xcorr’ and ‘corr’. (Figure 6D, E). All curve fitting was carried out using the MATLAB curve fitting toolbox.

#### Analysis of urethral sphincter EMG

For EMG data shown in Figures 5, artefacts of over 50 times the standard deviation for the recording were removed in MATLAB. The data was then RMS filtered using a 5 second moving window and smoothed using the MATLAB function ‘movmean’ over 1000 samples (a window of 0.32 s).

#### Analysis of non-voiding contractions (NVCs)

NVCs were identified in the intervoid periods only (Figure 7C). Pressure measurements during these periods were detrended (using the MATLAB function ’detrend’) and NVC peaks were detected using the MATLAB function ’findpeaks’, using parameters to select only those peaks greater than 0.1mmHg above the baseline, between 1.5 and 8 s wide, and at least 5 s from any other such peak. These provided well isolated examples of NVCs for analysis. Spike trains for each cell were binned (2s bin width for the longer timescales shown in Figure 7C; 0.5s bin width for Figure 7D) to extract estimates of firing rate around each NVC and to examine the difference in spike counts during 1.5s windows beginning 6s and 3s before each NVC. For shuffled data a vector of random times was generated for each recording (taken from intervoid periods only) and examined in the same way. The number of shuffled NVC times and real NVC times was equal in each recording.

#### Code availability

Custom MATLAB scripts used to analyse the data are available upon request.

## Supplemental Figure legends

**Supplemental Figure 1.**
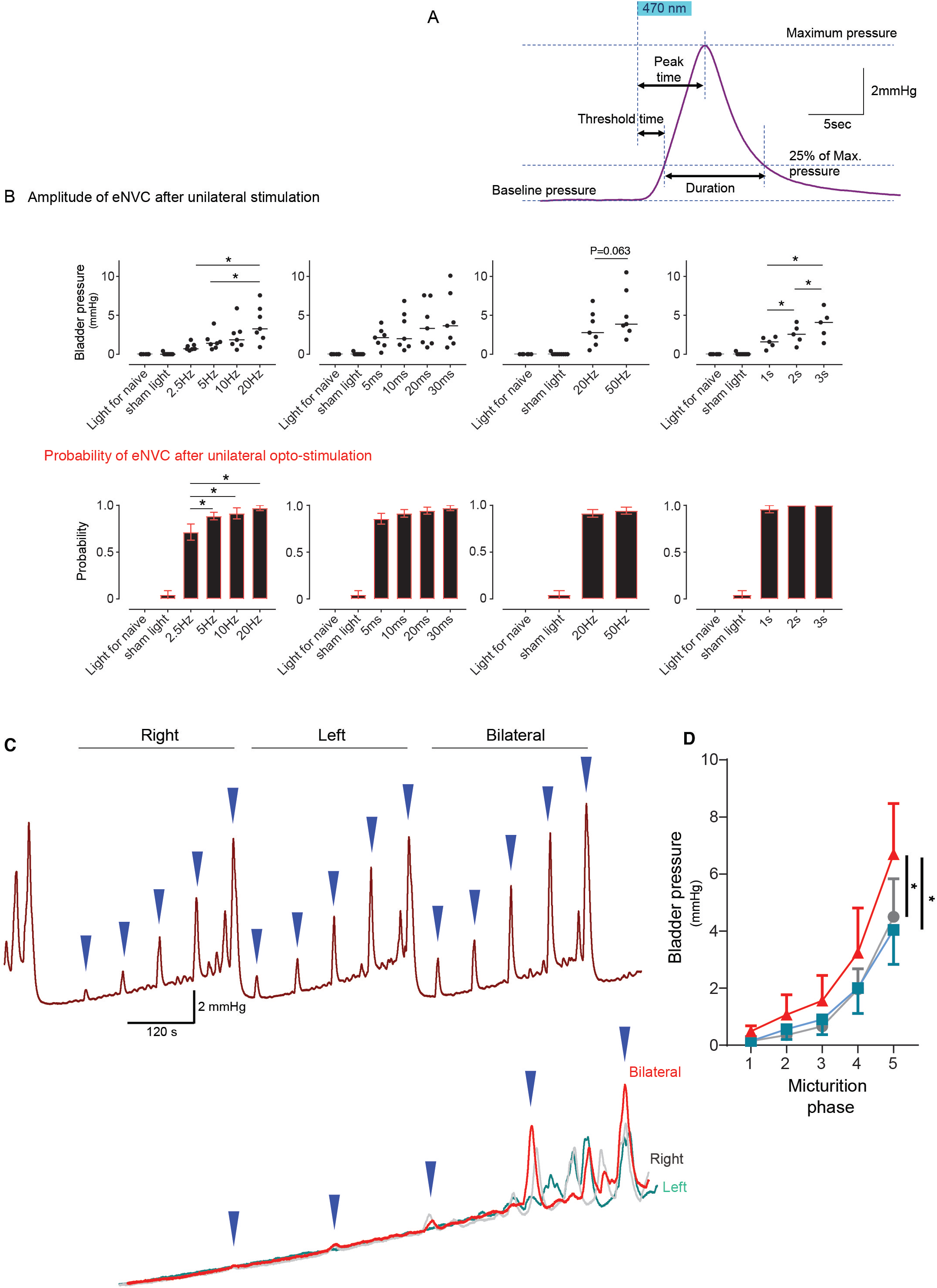
Characteristics of Barr^CRH^ evoked non-voiding contractions. A) Parameters of Barr^CRH^ evoked non-voiding contractions. Threshold calculated at 20% of the amplitude, duration was measured at the threshold pressure. The latency was taken as the time from start of stimulation for the pressure to reach threshold. B) The amplitude of eNVC increased with unilateral stimulation frequency (pulse length 20ms for 5s) but longer pulse durations (at 20Hz for 5 s) only tended to increase amplitude (*n=7* mice). Higher frequencies of stimulation (50Hz x10ms) did not substantially increase eNVC amplitude. Single longer continuous light pulses (1-3s) could also generate graded eNVCs. Similarly, the probability of generating an eNVC increased with stimulation frequency (pulse length 20ms for 5s) but pulse duration (frequency 20Hz for 5s) had little influence. Longer light pulses (1-3s) also reliably generated eNVCs. C) Comparison of unilateral with bilateral opto-activation of Barr^CRH^ neurons during continuous filling cystometry showing the graded increase in eNVC amplitude with phase of the micturition cycle and the augmented response to bilateral stimulation. D) Bilateral stimulation evoked larger eNVC, an effect that is more pronounced later in the micturition cycle (*n=7* mice).

**Supplemental Figure 2.**
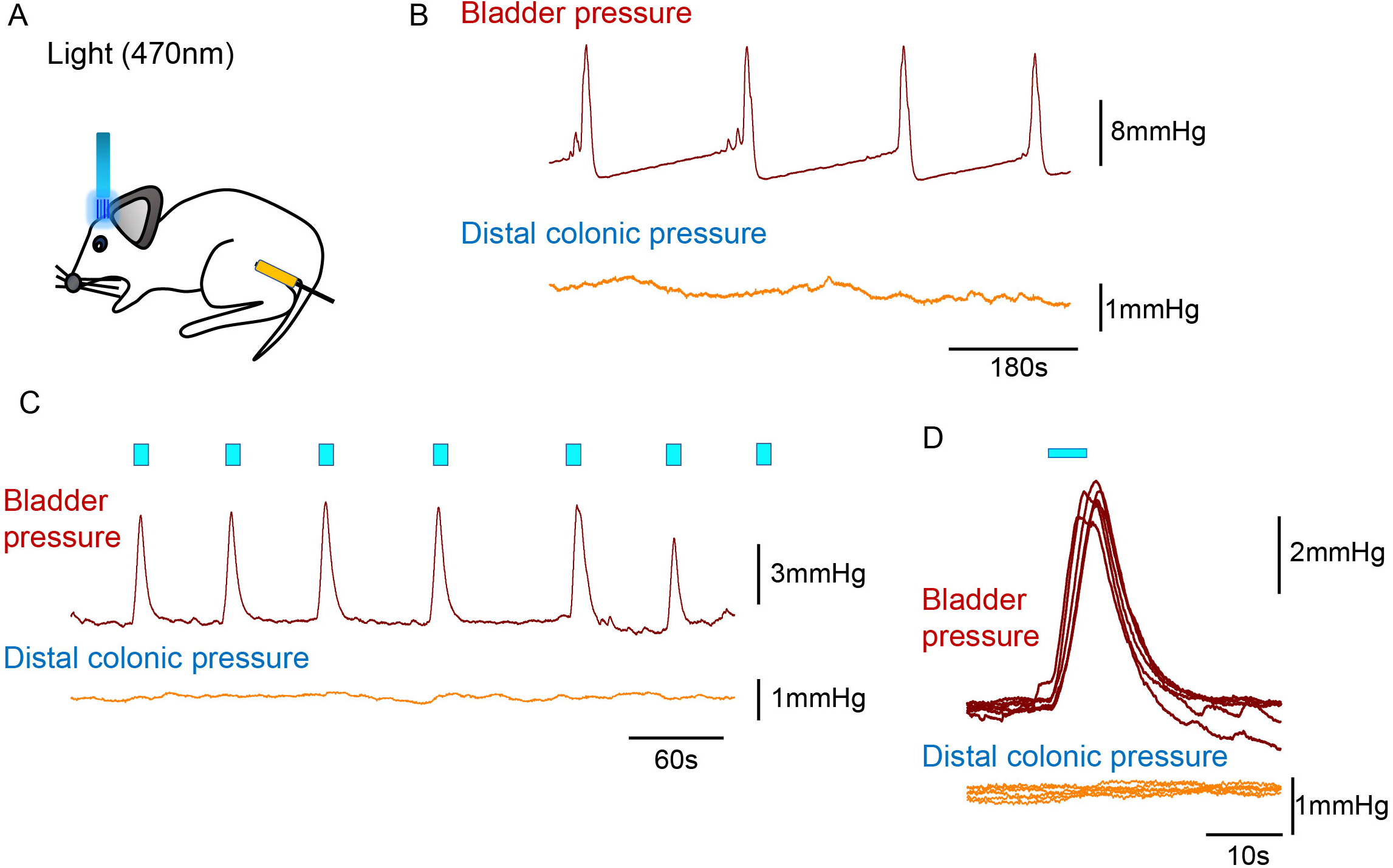
Barr^CRH^ neuronal activation does not cause contraction of the distal colon. Simultaneous recording of bladder pressure and rectal pressure with an intraluminal balloon showed no clear relationship between colonic pressure and the micturition cycle during continuous filling cystometry. Similarly, although opto-activation (20ms x 20Hz for 5s) of Barr^CRH^ generated eNVCs, there was no corresponding response in the distal colon.

**Supplemental Figure 3.**
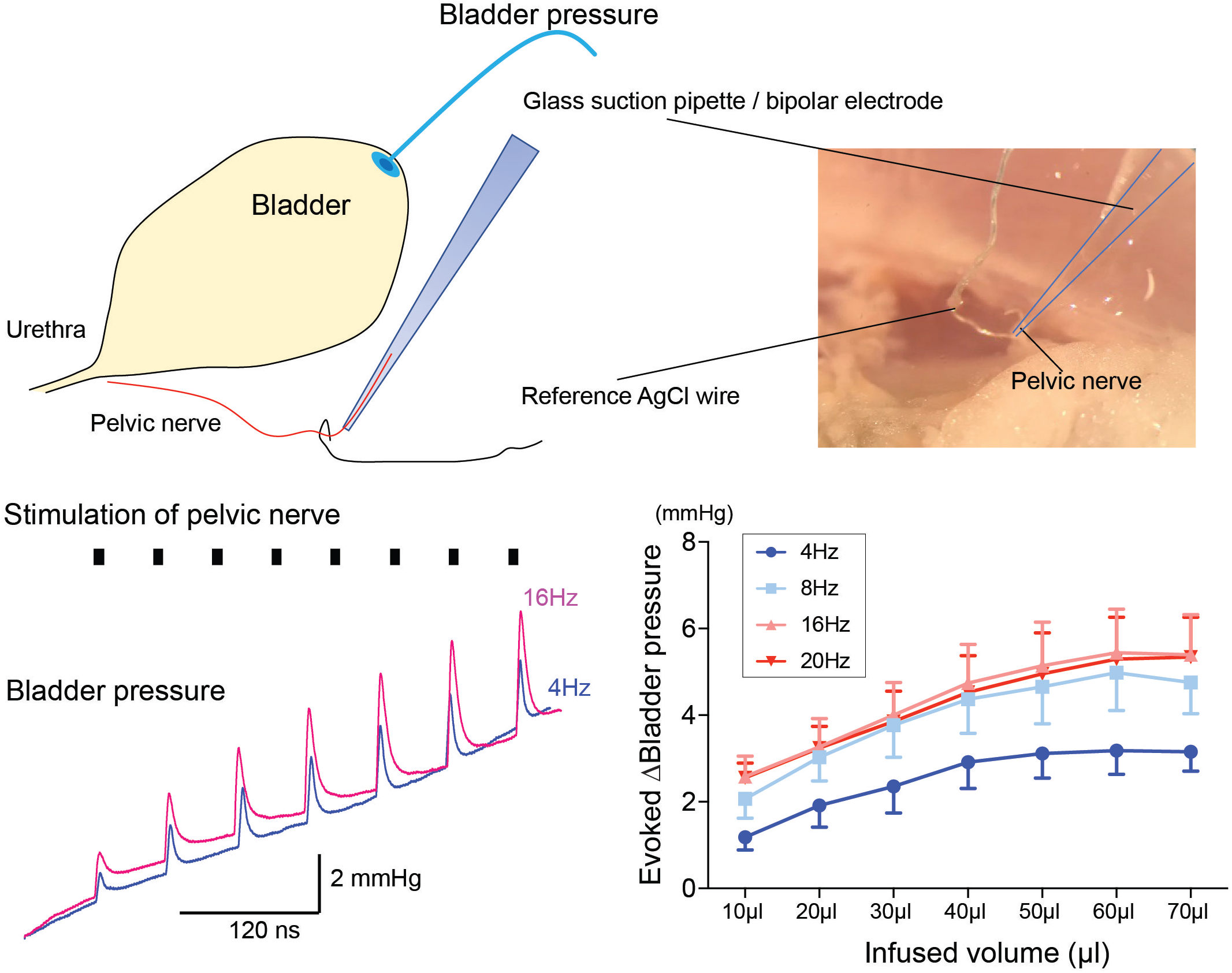
Dependence of pressure response to electrical stimulation of the pelvic nerve on bladder filling. Using the pithed decerebrate arterially perfused mouse preparation (*n=7*), the bladder pressure was monitored while the pelvic nerve was stimulated using a bipolar suction electrode (10V, 4-20Hz, train 3sec). Pelvic nerve stimulation evoked pressure responses that were dependent on stimulation frequency and on bladder distension.

**Supplemental Figure 4.**
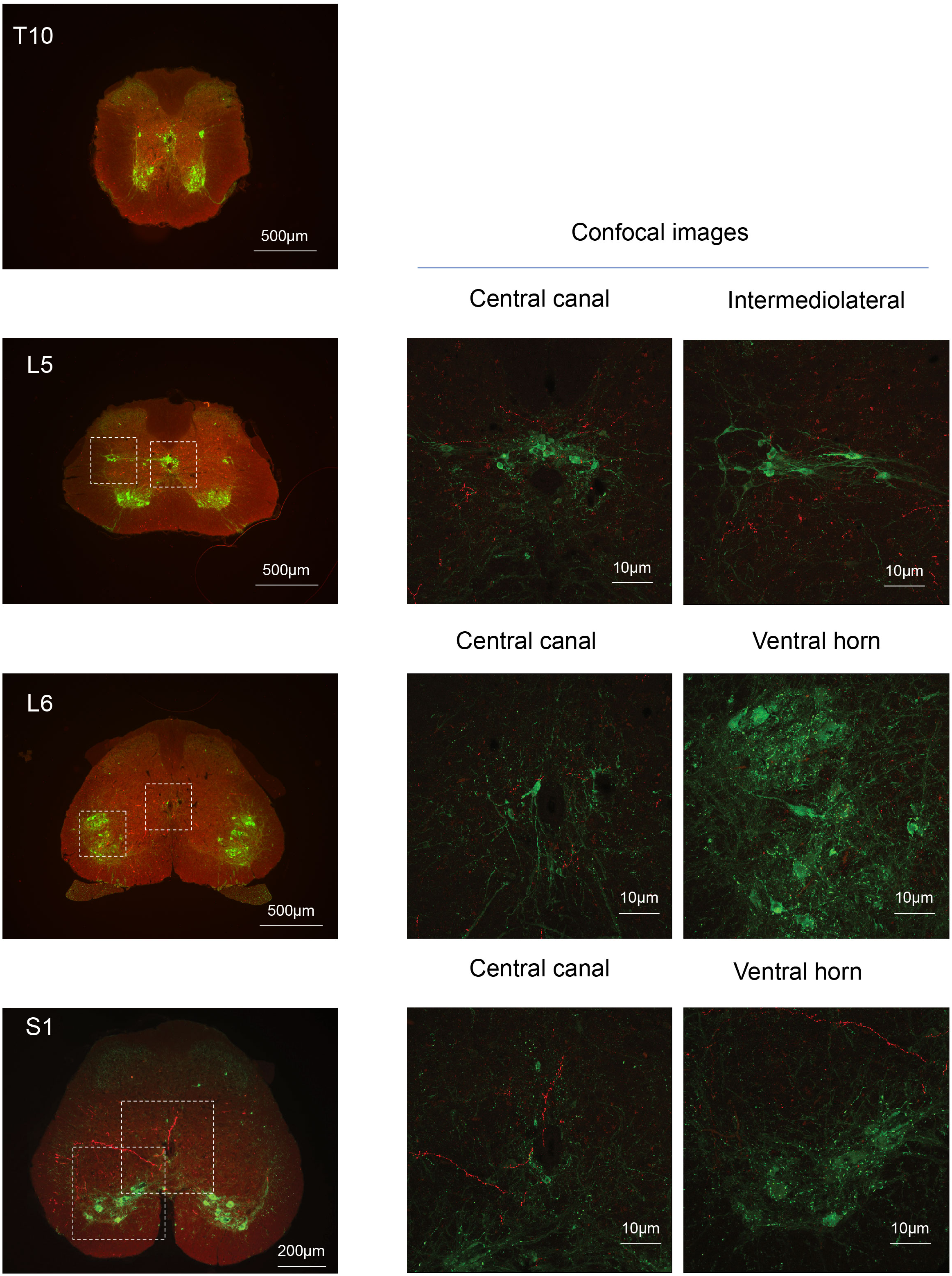
Spinal projections of Barr^CRH^ axons. Following unilateral transduction of Barr^CRH^ neurons with AAV-EF1*α*-DIO-ChR2-mCherry, transverse spinal cord sections (30µm) were cut from T10-S1 segments. Sections were processed for fluorescence immunocytochemistry for mCherry and Choline acetyltransferase to label filled Barr^CRH^ axons and somatic and autonomic motoneurons, respectively. These are represented as widefield and confocal images of blow-outs. The Barr^CRH^ axons show a lateralised distribution and can be seen to target the territory of parasympathetic preganglionic neurons at L5 and L6 as well as in the ventral horn area in S1. Note the absence of labelling close to somatic motoneurons and the sympathetic preganglionics at T10.

**Supplemental Figure 5.**
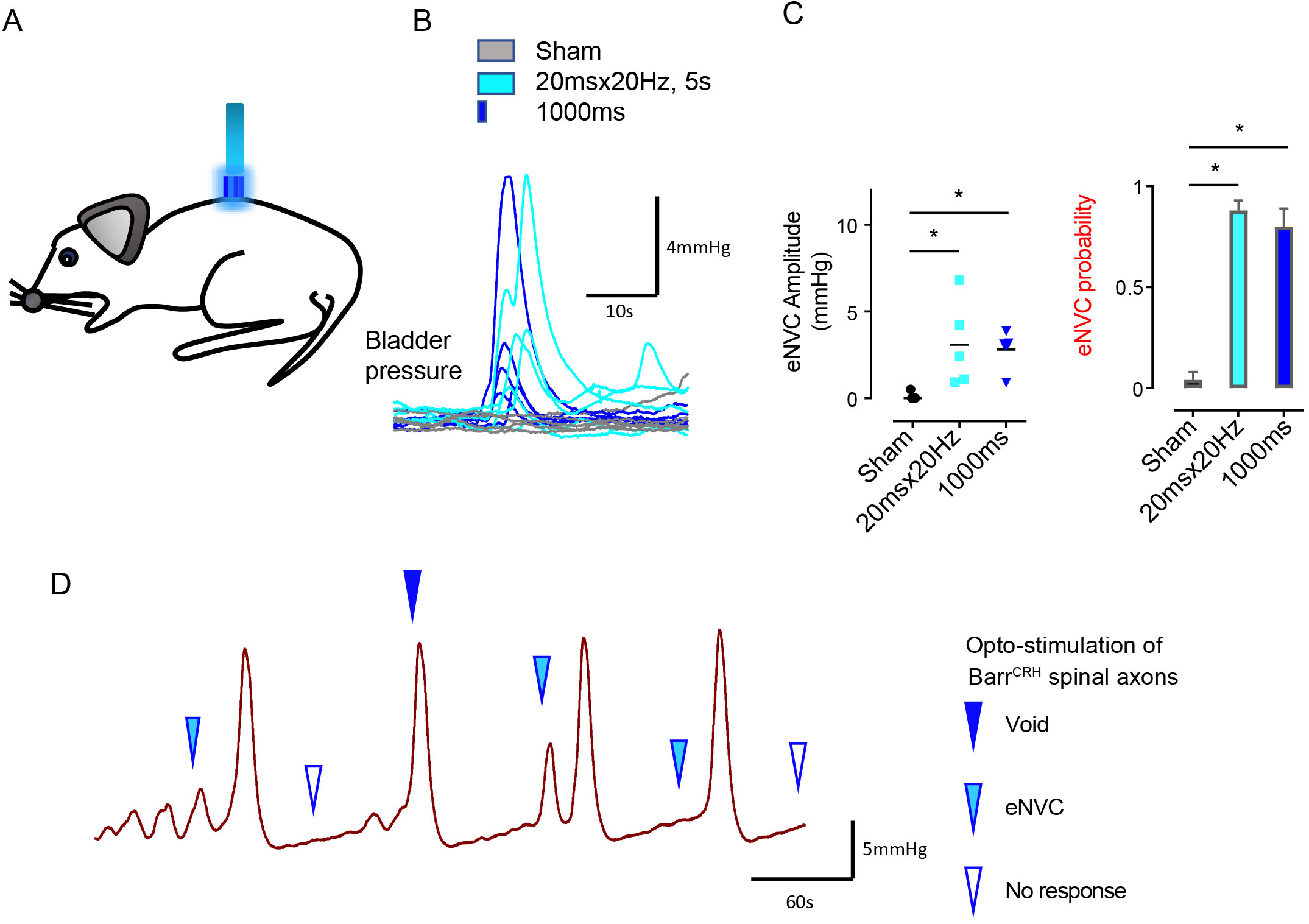
Spinal opto-activation of Barr^CRH^ axons generates eNVC and voids. A) The spinal cord was exposed at the level of T11-12 and illuminated from an optic fibre placed above the cord. B) Opto-activation (20Hz x 20ms for 5s or single 1s pulse) generated eNVCs (Related samples Friedman’s test by ranks). C) There was no difference in the eNVC in terms of amplitude or reliability between the two opto-stimulus patterns (n=5 mice). D) Opto-stimulation during continuous filling cystometry could generate full voiding contractions as well as eNVCs.

**Supplemental Figure 6.**
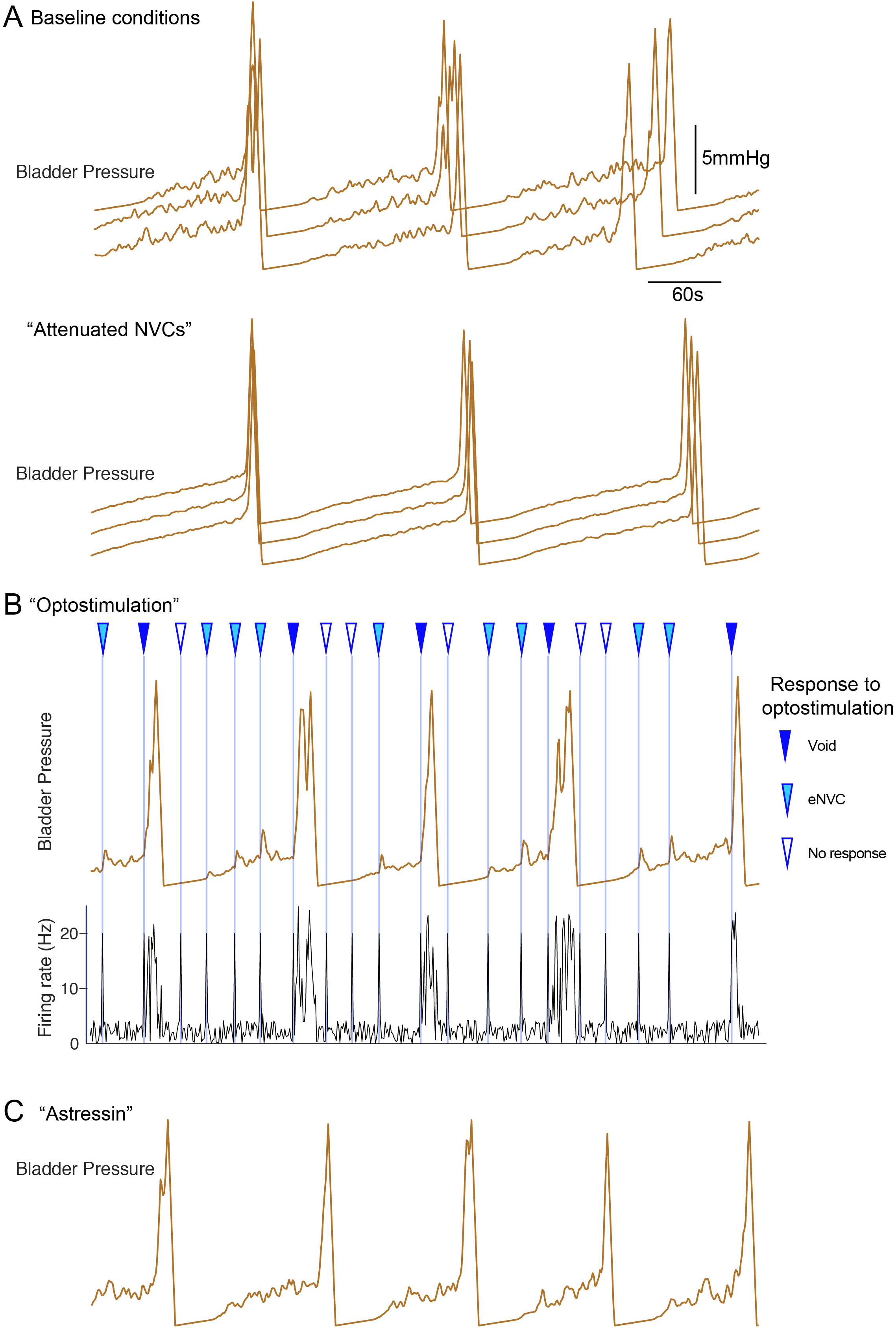
Inferential micturition model recapitulates observed behavior. A) Comparison of model outputs under basal conditions (upper panel, 3 consecutive runs) with a reduction in the variability in the BarrCRH firing (mean rates unchanged, lower panel). This attenuates the amplitude of the NVCs and delays the time to void indicating the importance of the NVCs in the micturition cycle. B) Simulation of optogenetic activation of BarrCRH neurons (20Hz x 1s) at different points in the micturition cycle (all other model parameters as the basal condition in A). This external drive increased micturition frequency by triggering voids (when stimuli fell later in the filling cycle). Note that opto-activation earlier in the filling cycle generated NVCs of varying amplitude and also could lead to ‘failures’ with no bladder contraction. C) A leftward shift in the mid-point (by 1hz) of the spinal modulation sigmoids mimicked the effect of CRH antagonist astressin by increasing the excitability of the spinal circuit. This increased the amplitude of NVCs and increased micturition frequency.

